# Intracellular screening of nanobodies reveals an intrabody that inhibits ITCH E3 ubiquitin ligase

**DOI:** 10.64898/2026.08.07.743588

**Authors:** Jeffrey W.T. Wang, Victor L. Lam, Ethan P. Dunn, Salvador Martinez, Rachel A. Jones, Anupama Sinha, Joshua T. Hong, Raga Krishnakumar, Joseph S. Schoeniger, Jennifer L. Schwedler, Christopher A. Sumner, Oscar A. Negrete, Steven S. Branda

## Abstract

Nanobodies are a class of small, monomeric camelid antibody fragments that can bind target antigens with high affinity and specificity. Their small size, structural simplicity, and limited reliance on disulfide bonding makes them attractive for intracellular expression for labeling and perturbing cellular processes in live cells. However, screening campaigns carried out exclusively *in vitro* often yield antigen binders that fail to perform well in live cells due to low expression, misfolding, or mistargeting. We demonstrate that traditional *in vitro* screening of a nanobody library combined with an intracellular bioluminescence resonant energy transfer (BRET) proximity sensor approach for sequence down-selection can yield strong *in vitro* binders that also perform well as intrabodies, in this case capable of binding to, and inhibiting the enzymatic activity of, ITCH E3 ubiquitin ligase in human cells. This strategy allows a more direct and scalable path toward intrabody discovery.

## Introduction

Intrabodies are antibody-based affinity ligands that are developed to function inside of live cells, to enable control of target protein (antigen) abundance, localization, activity, and labeling.^1–4^ However, antibodies as a class evolved to function in the extracellular environment and therefore may not fold properly, remain soluble, or maintain affinity and specificity when expressed in live cells^5^. Efforts have been made to modify antibodies to improve compatibility with intracellular use. These include rational engineering of stabilized mutants^6^ and the use of machine learning for intrabody domestication.^7^ In commonly used workflows, synthetic nanobodies (Nbs) derived from the antigen-binding fragment (V_H_H) of camelid heavy chain only antibodies^8^ are developed against an antigen of interest using phage display. Candidate binders are then verified to bind to antigen *in vitro* (using ELISA or BLI assays, for example), and then expressed in cells without further modification.^1,6,9^ As small (∼15 kDa) monomeric proteins with limited intramolecular disulfide bonding, Nbs have proven to be excellent starting points for intrabody development.^5^ Furthermore, the Nb scaffold itself has been subject to protein engineering for intracellular stability.^9^ However, many Nbs modified in this manner fail to perform well as intrabodies, making iterative optimization necessary.^6,9^ For instance, past studies converting Nbs to intrabodies yielded a success rate of less than 50%, motivating earlier evaluation of candidate Nbs in cells during intrabody development.^6,9^

We hypothesized that intrabody development might be improved by following the initial phage-display screen (Nb library biopanning) with a secondary screen for antigen binding in live cells. This approach should enable identification of Nb sequences that are compatible with intracellular expression and capable of binding to intracellular antigen. Secondary screening for *in cellulo* antigen binding has the added advantage of being more generalizable than screens based on a biological phenotype that is caused by Nb binding. For example, detecting intracellular Nb binding based on relocation of the target protein is useful only if the target protein has a distinctive subcellular localization profile.^6,7,9^ Similarly, screening for Nb-mediated inhibition of enzymatic activity is useful only if the target protein possesses enzymatic activity that is easily detected in cells.^10^

In the present study we demonstrate the utility of this approach through a campaign to develop Nb-based intrabodies that recognize a model intracellular protein: ITCH, a HECT-type E3 ubiquitin ligase. We chose to focus on ITCH because of its importance in immunity^11^, viral particle release^12,13^, and neuropathies like Alzheimer’s disease^14^ and because there were no previous reports of high-affinity, specific inhibitors of ITCH enzymatic activity (a small molecule inhibitor, clomipramine, has been shown to be a low-affinity, promiscuous inhibitor of HECT-type E3 ligases)^15,16^ nor of Nbs that bind to ITCH. Thus, new intrabodies for inhibiting ITCH activity in living cells could serve as valuable tools for the research community.

To detect Nb binding to ITCH *in cellulo* we used an intracellular protein proximity sensor system called nanoBRET (**nano**luciferase **B**ioluminescence **R**esonant **E**nergy **T**ransfer). nanoBRET relies on Förster resonance energy transfer (FRET) between a bioluminescent energy donor, in this case nanoluciferase (nLuc), and an energy acceptor, typically a fluorescent protein (FP).^17^ Two potentially interacting proteins are co-expressed, each as a fusion protein to either the donor or acceptor. When a protein-protein interaction (PPI) causes the donor and acceptor to be brought together in close proximity (c.a. <10nm) the resulting energy transfer can be detected as a redshifted bioluminescence (BRET signal).^18^ A unique challenge to executing a screen for intracellular Nb/antigen PPI arises from the extensive expression optimization required to develop a semi-quantitative nanoBRET assay capable of detecting antigen binding amongst many candidate Nb sequences. This is further complicated by the lack of a readily available positive control because, as in the case for ITCH and for most intrabody development campaigns, there were no previously identified Nbs for the intended target.

We found that Nb library biopanning followed by nanoBRET-based secondary screening enabled us to identify anti-ITCH intrabodies that show robust expression, binding to ITCH, and inhibition of ITCH enzymatic activities in live human cells. To overcome the assay development hurdle we used a known binding partner of ITCH, the Z protein from lymphocytic choriomeningitis virus (LCMV)^19,20^, as a surrogate for Nb binding when optimizing the nanoBRET assay conditions. Many of the Nbs displaying robust binding to intracellular ITCH in the nanoBRET assay were found to similarly bind ITCH *in vitro*, and several of these Nbs also inhibited ITCH enzymatic activities *in vitro* and *in cellulo*. Together, these results demonstrate a direct and scalable workflow for identifying functional intrabodies, which by design could be generalized for development of intrabodies against a wide variety of intracellular targets.

## Results

### nanoBRET assay development

The first step in developing a quantitative nanoBRET assay is to optimize the tagging and expression conditions for the interacting proteins (in this case ITCH and the Nbs recovered through biopanning against it). However, we lacked a positive control for our optimization efforts because Nbs against ITCH had not been reported previously. Accordingly, for purposes of optimizing our nanoBRET assay we paired ITCH with a viral protein with which it is known to interact: the Z protein from the lymphocytic choriomeningitis virus (LCMV). Because the intermolecular distance between the acceptor and donor has a strong effect on the BRET signal, the choice of N-terminal or C-terminal tagging can change the signal output for a given nanoBRET assay (**Figure 1A**). Furthermore, there may be unforeseen changes in protein stability or localization depending on the tagging configuration.

**Figure 1.**
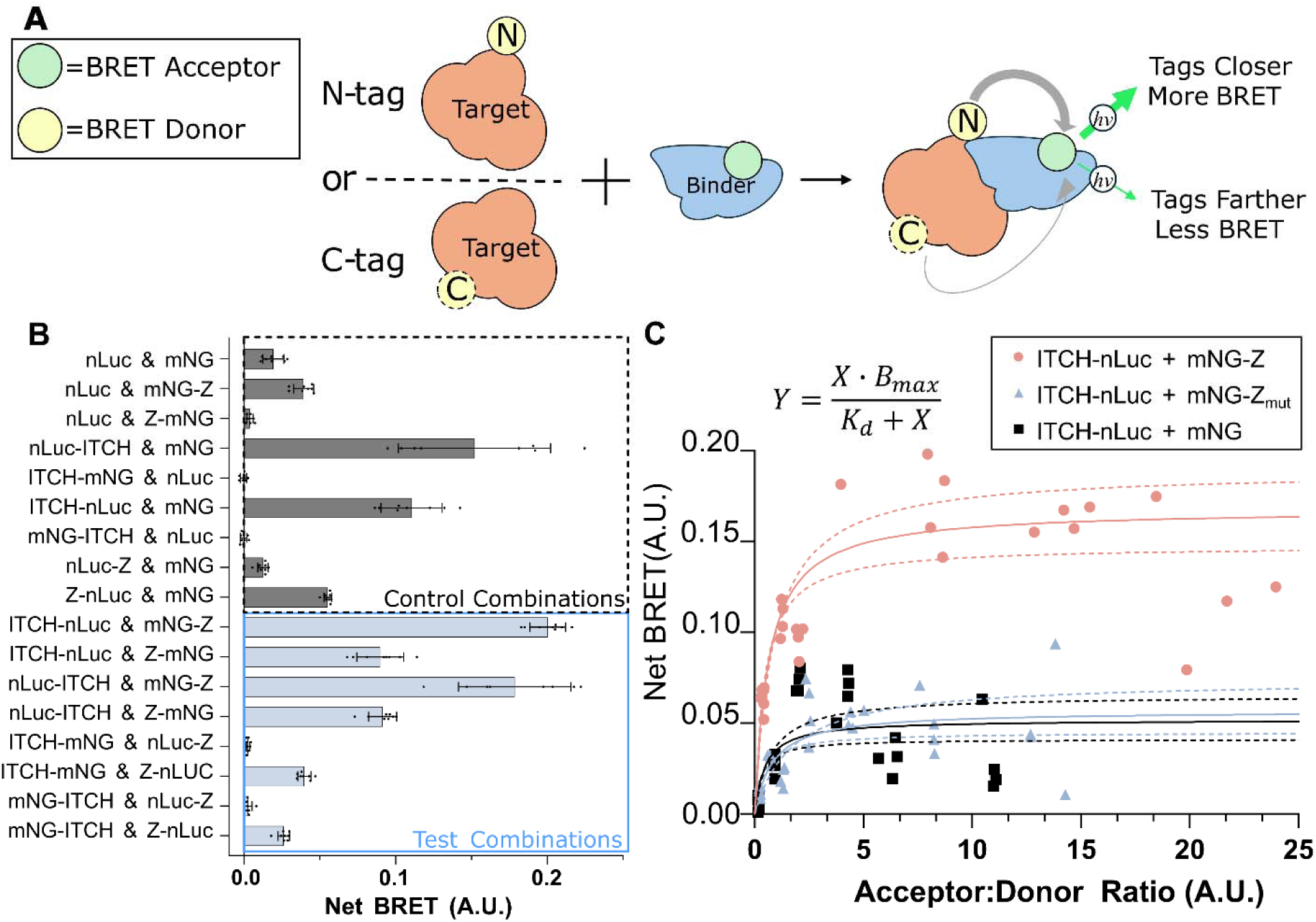
Development of a nanoBRET assay for detecting PPIs with ITCH. **A, Schematic illustrating the potential impact of tagging configuration on BRET signal.** The more closely the donor and acceptor are brought together, the higher the BRET efficiency. Accordingly, nanoBRET assays require optimization of the tagging configuration, as the choice of N-*versus* C-terminal fusion (nLuc-ITCH versus ITCH-nLuc) of the nanoBRET donor and acceptor moieties influences the maximum possible BRET signal achievable. **B, Optimizing tagging configuration for nanoBRET-mediated detection of intracellular PPI between ITCH and Z.** HEK293T cells in a 384-well plate were transfected with combinations of plasmids that express nLuc and mNG either alone or fused to ITCH or Z. Luminescence was then measured at 24 h post-transfection and net BRET signal was computed for each condition (8 replicate wells/condition). A.U. = arbitrary units. **C, nanoBRET plasmid titration assay to optimize detection of ITCH/Z interaction.** HEK293T cells were transfected with ITCH-nLuc and one of either mNG-Z, mNG-Z_mut_, or mNG at varying ratios. At 24 h post-transfection the cells were imaged for mNG expression and their BRET luminescence was measured. Net BRET was plotted against the relative quantity of donor and acceptor as computed by taking the ratio of mNG fluorescence intensity to total luminescence. The displayed single-site binding equilibrium model was fitted to the data, with the solid line representing the average of 500 bootstrapped fit parameters, and the dotted lines representing the 95% confidence intervals of the fit parameters.

We therefore combinatorially transfected ITCH and Z as N- or C- terminal fusions to either the nLuc donor or the FP acceptor. We used mNeonGreen (mNG) as the FP acceptor due to its reported higher efficiency relative to other FP acceptors^21^ and because in our hands an mNG-nLuc fusion exhibited higher BRET efficiency than an hfYFP-nLuc^22^ fusion (**Supplemental Figure 1**). Upon combinatorially transfecting ITCH and Z in each possible tagging orientation we found that the ITCH-nLuc and mNG-Z combination produced the highest BRET signal (**Figure 1B**). However, we observed comparable signal from a control condition: ITCH-nLuc combined with unfused mNG. This surprising result suggested that significant molecular crowding or aggregation was occurring under the transfection conditions used, leading to production of false-positive BRET signals. Such an effect might be amplified by ITCH’s recruitment to aggregating proteins.^23^ To investigate these issues we performed a similar set of combinatorial transfections that featured expression of a mutant Z protein (Z_mut_) in which the ITCH-binding PPXY interaction motif^19^ is replaced with alanines, and found that these yielded similar results, with most signal generated when the nLuc donor and mNG acceptor are fused to ITCH and Z_mut_ respectively (**Supplemental Figure 2**). These results indicate that overexpression of the nanoBRET constructs, leading to intracellular molecular crowding, was likely the primary source of false positive BRET signals.

Therefore, we carried out a nanoBRET acceptor saturation experiment using the configuration that produced the highest signal (ITCH-nLuc and mNG-Z) to identify transfection conditions that enable discrimination between a specific ITCH/Z interaction *versus* a nonspecific nLuc/mNG interaction, a key requirement for identifying anti-ITCH (α-ITCH) intrabodies. We transfected varying ratios of nLuc- and mNG-expressing plasmids to maximize the difference in BRET signal between the positive ITCH/Z interaction *versus* the negative Z_mut_ and mNG-only controls. Using the donor luminescence and acceptor fluorescence intensities as analogs for donor and acceptor abundance^24^ we fit a single-site equilibrium model to the data and calculated the 95% confidence intervals (95CI) around the modeled BRET signal as a function of acceptor:donor ratio. We observed higher BRET signal from the ITCH/Z pair as compared to the negative controls at acceptor:donor ratios greater than 5, with broad 95CI separation between positive and negative groups (**Figure 1C**). Importantly, we also observed a plateau in BRET signal that indicates a region of stable signal strength in which variations in expression level should have little effect on BRET signal intensities. Taken together, these results indicate that use of acceptor:donor ratios greater than 5 should support sensitive detection of ITCH PPIs while also avoiding false positive BRET signals due to intracellular molecular crowding.

### Intracellular Nb screening using nanoBRET

Having defined the conditions for detecting intracellular ITCH PPIs we used our nanoBRET assay system to screen Nbs for binding to ITCH *in cellulo*. Biopanning our synthetic Nb library^25^ had enabled us to enrich for Nbs that bind to the HECT domain of ITCH *in vitro* (see Methods section for details). The sequences encoding the 69 α-ITCH Nbs most frequently recovered in biopanning were individually cloned into an expression vector that directs genetic fusion to C-terminal mNG. We used this tagging configuration, rather than the N-terminal mNG tagging that was optimal for Z (see **Figure 1B**), in order to avoid potential steric clashes between the N-terminal tags and the complementarity determining region (CDR) loops of the Nbs.^26^ Each α-ITCH Nb-mNG fusion construct was then combined with the ITCH-nLuc fusion construct in concentrations expected to generate an acceptor:donor ratio falling within the previously determined operating plateau (**Figure 1C**). In parallel transfections we included mNG-Z as a positive control, mNG as a negative control, and Nb-mNG fusion constructs for two previously described off-target intrabodies (α-Histone^27^ and α-Actin^28^) as additional negative controls. Consistent with our previous results (**Figure 1B & C**), combining ITCH-nLuc with the mNG-Z positive control produced robust BRET signals in our nanoBRET assay (**Figure 2**). We also observed that combining ITCH-nLuc with each of several α-ITCH Nb sequences produced BRET signals similar in magnitude to those produced by the positive control. Note that the nanoBRET assay displayed unexpectedly high variance in BRET signal (**Figure 2**) as compared to previous nanoBRET experiments (**Figure 1**) and benchmarks established for our robotic liquid handling transfection procedure (**Supplemental Figure 3**). This variance may have originated from unexpected interplay between ITCH and the α-ITCH Nbs, or from a run-specific systematic robotic liquid handling error. In any case, the high variance led us to select for downstream analysis the α-ITCH Nbs with the top seven and the 10^th^ highest BRET signal, rather than use a statistically-defined cutoff. Therefore, a total of eight α-ITCH Nbs that displayed strong, reproducible interaction with ITCH in cells, as indicated by nanoBRET assay measurements, were selected for further analysis. These α-ITCH intrabodies are listed in **Table 1**.

**Figure 2.**
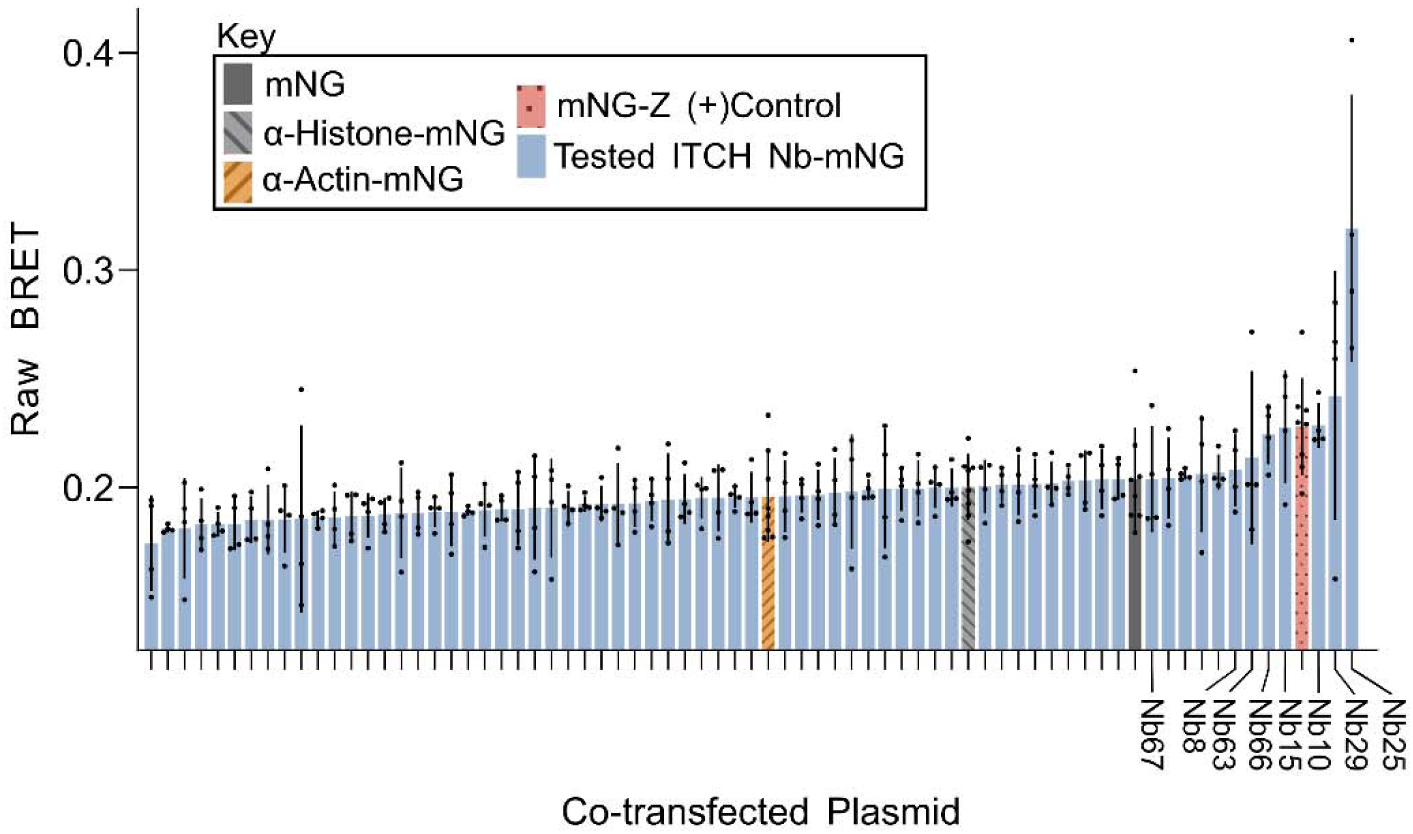
Intracellular screening of α-ITCH Nbs using nanoBRET. HEK293T cells were co-transfected with the CHCH fusion construct as well as either an α-ITCH Nb-mNG fusion construct, a positive control fusion construct (mNG-Z), a negative control fusion construct (α-Actin or α-Histone), or an mNG expression construct (an additional negative control). Each nanoBRET assay was carried out in either four replicate wells (α-ITCH Nb constructs) or eight replicate wells (control constructs), with raw BRET signal measured. 69 α-ITCH Nbs were individually tested for PPI with intracellular ITCH. For each PPI condition tested (X-axis) the mean raw BRET signal was measured (Y-axis). Black dots indicate raw BRET signal in individual wells and black lines indicate standard deviation. Candidate α-ITCH Nb sequences subsequently purified for *in vitro* validation are labeled.

**Table 1.**
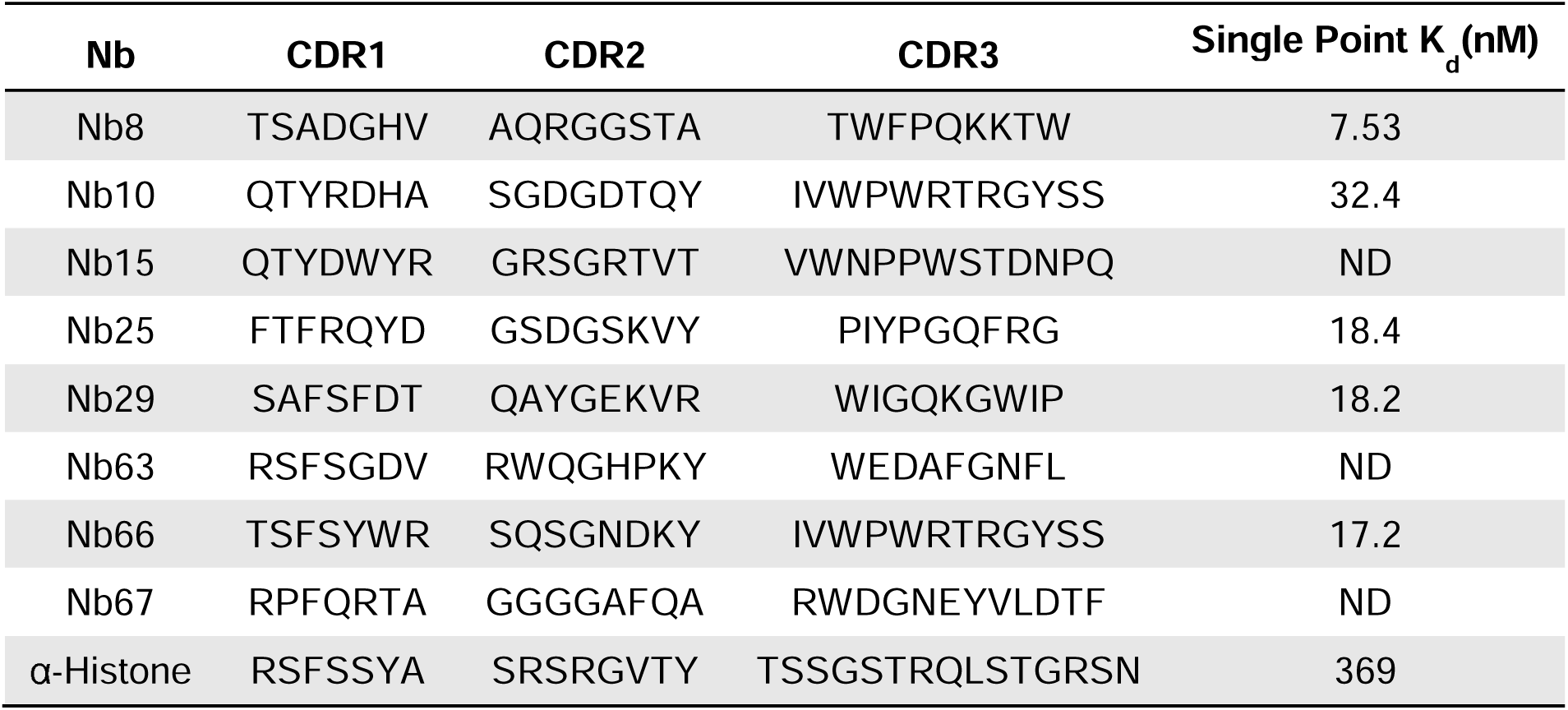
α-ITCH intrabodies selected for further testing. For each intrabody the three CDR sequences, as well as the single-point K_d_ value for binding to the ITCH HECT domain *in vitro*, are shown. The negative control α-Histone intrabody is included for comparison. ND: Not determined.

### α-ITCH intrabodies bind to ITCH and inhibit its enzymatic function *in vitro*

Sequences encoding the eight α-ITCH intrabodies selected for further analysis were individually cloned into an expression vector for production by *Escherichia coli* to support *in vitro* studies. However, only five of the α-ITCH intrabodies were produced in quantities sufficient for high-confidence measurement of binding to the HECT domain of ITCH *in vitro* as assessed *via* Bio-Layer Interferometry (BLI) analysis (**Figure 3A** and **Table 1**). For these five α-ITCH intrabodies the measured K_d_ values ranged from 7.53-32.4 nM, whereas the negative control α-Histone intrabody displayed an order of magnitude weaker K_d_ (369 nM). Notably, the low-yield preparations of the other three α-ITCH intrabodies (Nb15, Nb8, and Nb67) did display detectable binding to the ITCH HECT domain (**Supplemental Figure 4**). In summary, these results indicate that α-ITCH intrabodies identified using our nanoBRET assay also displayed high-affinity binding to the ITCH HECT domain *in vitro*.

**Figure 3.**
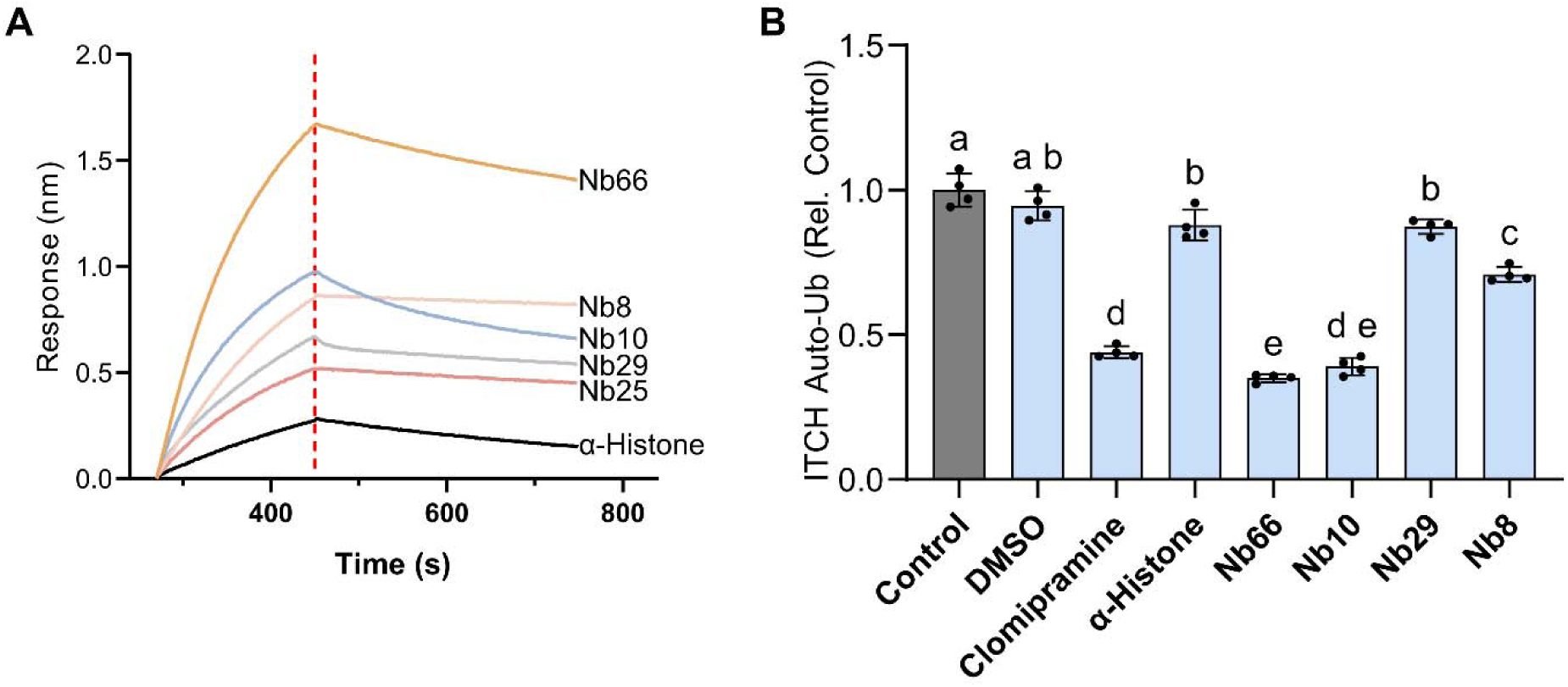
α-ITCH intrabodies tested for binding to ITCH and inhibition of its auto-ubiquitination activity *in vitro*. **A,** α**-ITCH intrabody binding to the ITCH HECT domain *in vitro* as measured *via* biolayer interferometry (BLI).** The BLI sensor was loaded with a fixed concentration of Nb (62.5nM) and then pulsed with ITCH HECT domain (250 nM) for 450 seconds. The dashed red line indicates the transition from Nb-ITCH association to dissociation. **B,** α**-ITCH intrabody mediated inhibition of ITCH auto-ubiquitination *in vitro* as measured *via* TR-FRET.** Each Nb (200 nM) was added to the ITCH HECT domain (100 nm) to achieve two-fold molar excess, and ITCH auto-ubiquitination was measured using an end-point TR-FRET assay (n=4 replicate wells).

Given that the HECT domain is the catalytic core of ITCH, we wondered whether the α-ITCH intrabodies that bind to it with high affinity might inhibit ITCH’s E3 ubiquitination activity. As an initial step in addressing this question we tested several of the α-ITCH intrabodies for ability to inhibit ITCH auto-ubiquitination as detected using a time-resolved FRET (TR-FRET) based assay system. In these experiments ITCH was incubated with an E1 enzyme, an E2 enzyme, and a mixture of ubiquitin (Ub) labeled with either terbium (Tb; donor) or fluorescein (FITC; acceptor). ITCH auto-ubiquitination activity generates a polyubiquitin chain on ITCH that can be detected based on FRET between adjacent Ub-Tb and Ub-FITC molecules. Using this assay, we determined that α-ITCH intrabodies Nb10 and Nb66, when individually administered in two-fold stoichiometric excess, inhibited ITCH auto-ubiquitination activity by ∼60% (**Figure 3B**). This degree of inhibition was comparable to that achieved through addition of clomipramine (250 μM), a potent though non-specific inhibitor of ITCH.^15,16^ The full kinetic traces for the TR-FRET experiment are shown in **Supplemental Figure 5**. It is notable that Nb10 and Nb66 possess the same CDR3 sequence (**Table 1**), suggesting that they may bind to a shared epitope on ITCH and thereby similarly inhibit its enzymatic activity.

Single-point BLI measurements indicated that Nb66 displayed higher affinity binding to ITCH as compared to Nb10, so we proceeded with further analysis of Nb66. We determined the average K_d_ of Nb66 at various concentrations of the ITCH HECT domain to be 7.2 nM ±3.7 nM (**Supplemental Figure 6)** and verified Nb66 binding to full-length ITCH via BLI (**Supplemental Figure 7**).

We then investigated whether Nb66 was capable of inhibiting ITCH poly-ubiquitination as well as trans-ubiquitination of a known target substrate (LATS1)^29^ *in vitro*. In these experiments full-length ITCH was combined with FLAG-tagged LATS1 (FLAG-LATS1), HA-tagged ubiquitin (HA-Ub), a ubiquitin-activating enzyme (UbE1), and a ubiquitin-conjugating enzyme (UbE2D4). Nb66 was then added to the reaction in three-fold molar excess as compared to ITCH. After addition of ATP and incubation at 37`°` C for 1.5 h the reaction was quenched and analyzed via Western blot. Detection of HA-Ub using an α-HA antibody revealed that ITCH mediated assembly of polyubiquitin chains, as evident from accumulation of high-molecular weight products; and that Nb66 inhibited this activity, as evident from reduced levels of these high-molecular weight products, comparable to those observed in reactions lacking ITCH (**Figure 4A**). Additionally, ITCH-mediated poly-ubiquitination of FLAG-LATS1 can be inferred from reduced intensity of a band at ∼140 kDa (the expected molecular weight of FLAG-LATS1); and Nb66-mediated inhibition of this activity can be inferred from increased intensity of this band, comparable to that observed in reactions lacking ITCH. Direct detection of FLAG-LATS1 using an α-FLAG antibody confirmed that ITCH mediated trans-ubiquitination of FLAG-LATS1, as evident from accumulation of high-molecular weight products and concomitant loss of the ∼140 kDa band; and that Nb66 inhibited this activity, as evident from the converse trends (**Figure 4B**). Taken together, these results indicate that Nb66 effectively inhibits ITCH auto-ubiquitination, poly-ubiquitination, and trans-ubiquitination activities *in vitro*.

**Figure 4.**
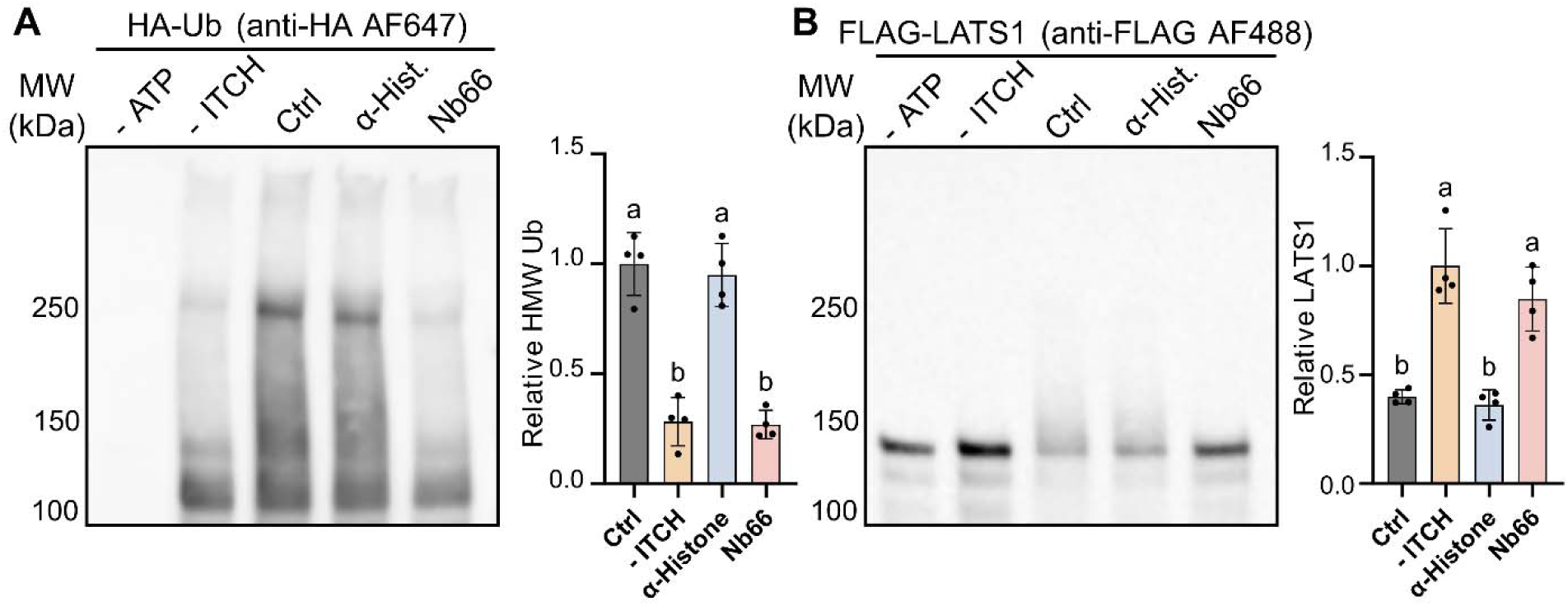
α-ITCH intrabody Nb66 tested for inhibition of ITCH poly-ubiquitination and trans-ubiquitination activities *in vitro*. Full-length ITCH (0.6 µM) was combined with affinity-tagged target substrate LATS1 (FLAG-LATS1; 1 µM), affinity-tagged ubiquitin (HA-Ub; 75 µM), and a ubiquitin-activating enzyme (UbE1; 2 µM) and ubiquitin-conjugating enzyme (UbE2D4; 8 µM). Where specified, an α-ITCH (Nb66) or negative control (α-Histone) intrabody was added to the reaction (1.8 µM); alternatively, only buffer was added (Ctrl). Upon addition of ATP (2 mM) the reaction was incubated at 37`°` C for 1.5 h and analyzed via Western blot. **A,** α**-ITCH intrabody mediated inhibition of ITCH poly-ubiquitination activity.** Reaction products bearing HA-Ub were detected using an α-HA antibody labeled with AlexaFluor 647 (AF647) (left). Each bar indicates the average intensity of the poly-HA-Ub smear normalized to that of the buffer-treatment negative control reaction (Ctrl), and each line indicates standard deviation (n=4 replicates *per* reaction type) (right). A one-way ANOVA with Tukey’s t-test with multiple comparisons correction was performed; groups that did not differ to a significant degree (p>0.05) are marked with the same letter, whereas those that did differ to a significant degree (p<0.001) are marked with different letters. **B,** α**-ITCH intrabody mediated inhibition of ITCH trans-ubiquitination activity.** FLAG-LATS1 was detected using an α-FLAG antibody labeled with AF488 (left). ITCH-mediated poly-ubiquitination of FLAG-LATS1 (∼140kDa) produces a higher-molecular weight smear with concomitant disappearance of the ∼140 kDa band. Each bar indicates the average intensity of the ∼140 kDa FLAG-LATS1 band normalized to that of the no-ITCH negative control reaction (-ITCH), and each line indicates standard deviation (n=4 replicates *per* reaction type) (right). A one-way ANOVA with Tukey’s t-test with multiple comparisons correction was performed; groups that did not differ to a significant degree (p>0.05) are marked with the same letter, whereas those that did differ to a significant degree (p<0.001) are marked with different letters.

In positive control reactions, clomipramine (a small-molecule inhibitor of ITCH) was added (250 μM) instead of a Nb; and in negative control reactions only buffered saline was added. TR-FRET signal from each reaction was normalized to a photobleaching control and then to the average of the positive control reactions. A one-way ANOVA with Tukey’s multiple comparisons test between the means of each treatment was performed. Treatments that did not have significantly different effects on TR-FRET signal (p>0.05) are marked with the same letter (e.g., both a, as for Control *versus* DMSO), whereas those that did have significantly different effects (p<0.05) are marked with different letters (e.g., a *versus* b, as for Control *versus* α-Histone).

### Nb66 binds and inhibits ITCH in human cells

Given that Nb66 displayed high-affinity binding to ITCH and inhibition of its enzymatic activity *in vitro* we sought to characterize its interactions with ITCH in live human cells. To assess Nb66-ITCH PPIs *in cellulo* we introduced a doxycycline-inducible Nb66-mNG expression construct into HEK293T cells, and transfected the stable cell line with an mScarlet-ITCH expression construct. We found that when expressed alone Nb66-mNG is uniformly distributed throughout the cell, whereas co-expression with mScarlet-ITCH results in exclusion of Nb66-mNG from the nucleus, matching the subcellular distribution of mScarlet-ITCH itself (**Figure 5A**). We also found that transfection of the Nb66-mNG stable line with an nLuc-ITCH expression construct generated a 17-fold increase in BRET signal as compared to a matched control (transfection of a α-Histone-mNG stable line with the nLuc-ITCH construct) (**Figure 5B**). This result strongly suggests that Nb66 binds to ITCH *in cellulo* and corroborates the results from our nanoBRET screening experiments (**Figure 2**). We further confirmed this intracellular PPI using co-immunoprecipitation (co-IP) experiments. We found that pull-down of Nb66-mNG (using mNG as the IP handle) enabled co-IP of HA-ITCH (as detected using α-HA antibody in Western analysis) (**Figure 5C**). In contrast, pull-down from cells expressing mNG alone, or expressing α-Histone-mNG, failed to co-IP HA-ITCH, as expected; however, it should be noted that pull-down of α-Histone-mNG was inefficient, likely because much of it partitioned into an insoluble fraction of the cell lysate (**Supplemental Figure 8**). Taken together, the results from these three orthogonal series of experiments confirm that the Nb66 intrabody binds to ITCH in human cells, validating our nanoBRET screening results (**Figure 2**).

**Figure 5.**
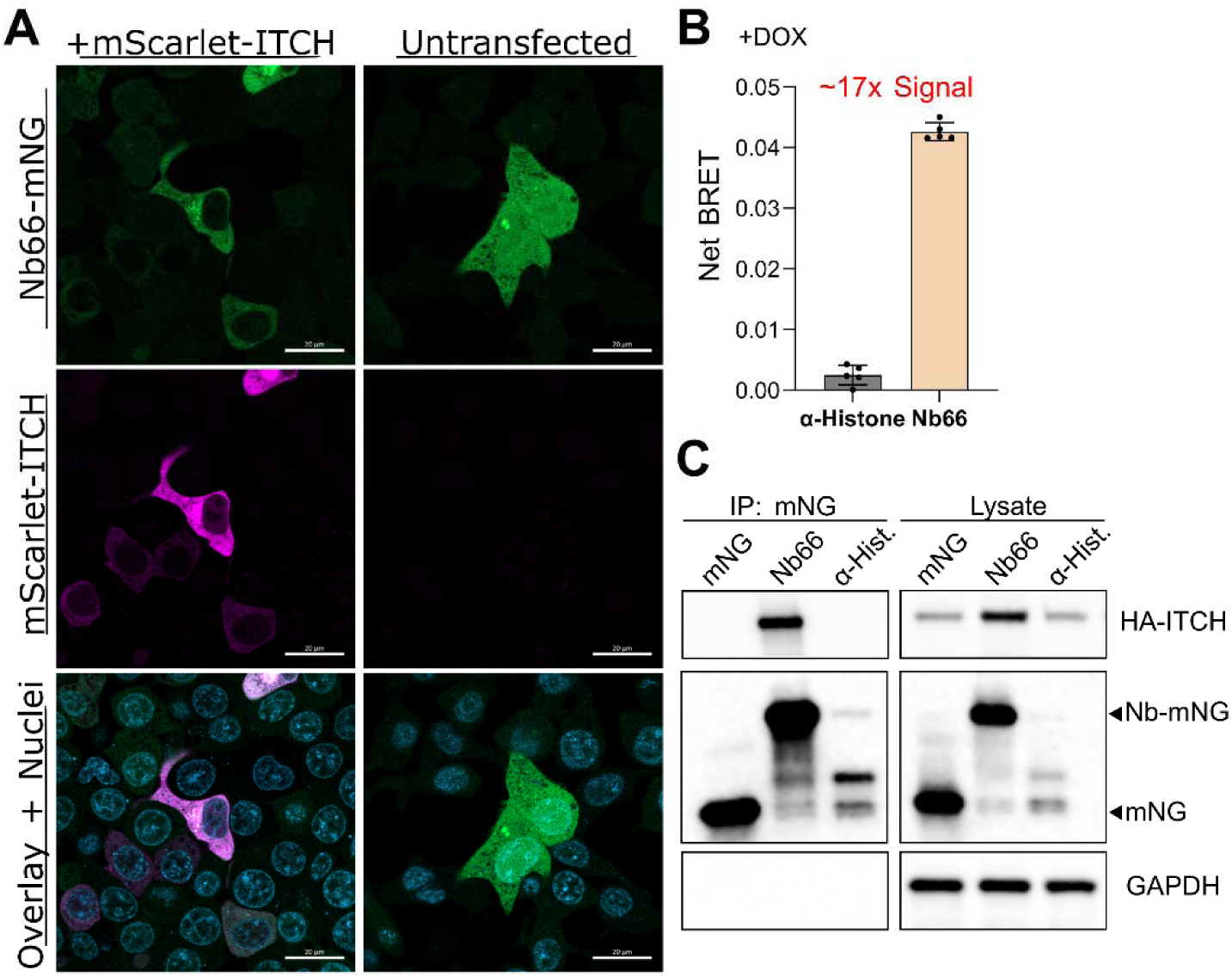
Validation of Nb66 binding to ITCH in human cells. **A, Colocalization analysis of Nb66 and ITCH.** HEK293T cells with stable integration of an expression construct for doxycycline-inducible production of the Nb66-mNG fusion protein were transfected with an mScarlet-ITCH expression construct (left), or were not transfected (right), in the presence of doxycycline hyclate (500 ng/mL); and subcellular localization of Nb-mNG (green) and mScarlet-ITCH (magenta), as compared to Hoechst stained nuclei (blue), was analyzed *via* confocal microscopy. Scale bar: 20μm. **B, nanoBRET analysis of Nb66 binding to ITCH.** HEK293T cells with stable integration of an expression construct for doxycycline-inducible production of either Nb66-mNG or α-Histone-mNG were transfected with an ITCH-nLuc expression construct in the presence of doxycycline hyclate (500 ng/mL); and BRET signal was measured at 24 h post-transfection. Net BRET was computed using a correction factor obtained from wells transfected in the absence of doxycycline induction. **C, Co-immunoprecipitation analysis of Nb66 binding to ITCH.** HEK293T cells were transfected with an HA-ITCH expression construct in combination with an mNG, Nb66-mNG, or α-Histone-mNG expression construct. An α-mNG antibody was used for pull-down of mNG from the cell lysates, and both the immunoprecipitates (left) and original lysates (right) were subjected to Western analysis to detect HA-ITCH (α-HA antibody; top); mNG, Nb66-mNG, or α-Histone-mNG (α-mNG antibody; middle); and GAPDH (α-GAPDH antibody; bottom).

Finally, we investigated whether Nb66 is capable of inhibiting ITCH auto-ubiquitination and poly-ubiquitination activities in human cells. These experiments leveraged a previously described assay for measuring the auto-ubiquitination activity of intracellular ITCH.^30^ HEK293T cells were co-transfected with HA-ITCH and myc-Ub expression constructs in combination with the Nb66-mNG expression construct or a negative control (either pUC19 or the α-Histone-mNG expression construct). After inhibiting proteasomal degradation (to allow ubiquitinated proteins to accumulate in the cells) we pulled down HA-ITCH (using the α-HA antibody) and assessed the immunoprecipitates for ubiquitinated adducts *via* Western analysis (using the α-HA antibody to detect HA-ITCH itself and an α-myc antibody to detect myc-Ub moieties associated with it). ITCH auto-ubiquitination presents itself as a mass-shifted HA-ITCH and high molecular weight myc-Ub smear.^30^ As seen in **Figure 6A**, expression of Nb66-mNG resulted in significantly reduced levels of mono-ubiquitinated (Ub_1_) and di-ubiquitinated (Ub_2_) HA-ITCH adducts as compared to the levels observed in the pUC19 and α-Histone-mNG controls (p<0.0001), indicating that Nb66-mNG inhibited HA-ITCH auto-ubiquitination activity. Similarly, **Figure 6B** shows that expression of Nb66-mNG resulted in significantly reduced levels of myc-Ub in the HA-ITCH immunoprecipitates (p = 0.024), indicating that Nb66-mNG inhibited HA-ITCH poly-ubiquitination activity. These results in combination with binding validation (**Figure 5**) confirm that the Nb66 intrabody binds ITCH and inhibits its auto-ubiquitination and poly-ubiquitination activities in human cells.

**Figure 6.**
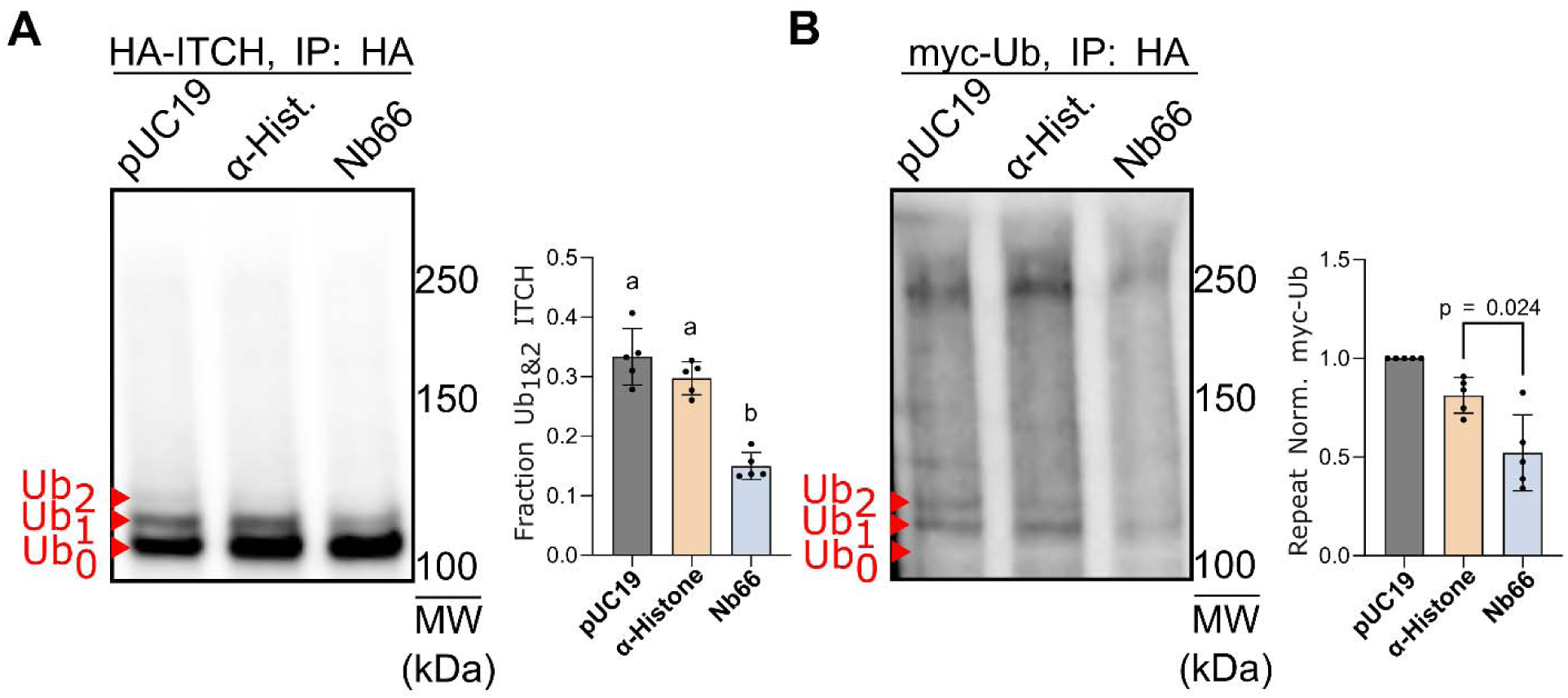
Nb66 tested for inhibition of ITCH auto-ubiquitination and poly-ubiquitination activities in human cells. HEK293T cells were transfected with HA-ITCH and myc-Ub expression constructs in combination with pUC19, an α-Histone-mNG expression construct, or an Nb66-mNG expression construct. A proteasome inhibitor (MG-132) was added at 20 h post-transfection, and cell lysates were collected at 25 h post-transfection. Immunoprecipitation (IP) of HA-ITCH was accomplished using an α-HA antibody, and the recovered proteins were subjected to Western analysis using two different detection antibodies. **A, Nb66 inhibition of ITCH auto-ubiquitination.** HA-ITCH was detected using α-HA antibody, revealing unmodified HA-ITCH (Ub_0_) as well as ubiquitinated adducts (Ub_1_ and Ub_2_) in the HA-ITCH immunoprecipitates (left). Each form of HA-ITCH was quantified *via* densitometry analysis, and the proportion of HA-ITCH that was ubiquitinated (i.e., the ratio of Ub_1_ and Ub_2_ adducts relative to total HA-ITCH) was calculated; the mean (bar) and standard deviation (line) values from five replicate experiments are shown (right). A one-way ANOVA with Tukey’s t-test with multiple comparisons correction was performed; groups that did not differ to a significant degree (p>0.05) are marked with the same letter, whereas those that did differ to a significant degree (p<0.0001) are marked with different letters. **B, Nb66 inhibition of ITCH poly-ubiquitination.** myc-Ub was detected using an α-myc antibody, revealing unmodified proteins (Ub_0_) and ubiquitinated adducts (e.g., Ub_1_ and Ub_2_) in the HA-ITCH immunoprecipitates (left). Densitometry analysis was carried out, and the proportion of poly-ubiquitinated HA-ITCH (intensity of the myc-Ub smear) in total HA-ITCH (intensity of HA signal in the same sample, as detected in Fig. 6A) was calculated, with repeat-to-repeat variation corrected by normalizing to the pUC19 control within each repeat; the mean (bar) and standard deviation (line) values from five replicate experiments are shown (right). Welch’s t-test was performed between Nb66 and α-Histone groups (p = 0.024).

## Discussion

We sought to address the challenge of developing intrabodies by establishing a workflow in which Nb library biopanning is followed by a nanoBRET-based secondary screen for Nb expression and target binding in live human cells. This approach enabled us to directly test intracellular performance (both expression and target binding) at moderately high throughput (69 Nbs tested in parallel) (**Figure 2**) without need for prior down-selection based on *in vitro* performance (target binding, as assessed *via* ELISA or BLI for example). In fact, we found that α-ITCH Nbs displaying robust expression and target binding *in cellulo* (**Figure 2**, **Figure 5**) generally also displayed robust target binding *in vitro* (**Figure 3A**, **Table 1**). Moreover, we found that several of the high-performing α-ITCH intrabodies were capable of inhibiting ITCH’s enzymatic activities *in vitro* (**Figure 3B**, **Figure 4**), and at least one (Nb66) also inhibited these activities *in cellulo* (**Figure 6**).

Importantly, our approach should be generalizable to support development of Nbs against a wide variety of intracellular targets. Mounting a new intrabody development campaign requires outfitting the Nbs and target with nanoBRET donor and acceptor moieties, but this is a relatively straightforward process that can leverage modular cloning techniques^31^ and ready-made expression plasmids.^17^ that a binding partner for the target is required for optimization of the nanoBRET assay but may not be available, as when natural PPI partners are not known and target-specific Nbs have not been developed yet. We propose that this problem could be addressed in future work by taking advantage of Nbs that have been shown to bind to epitope tags *in cellulo*, such as those recognizing V5^32^ and ALFA^33^; with one simple additional step (fusion of the epitope tag to the target protein) the needed binding partner would become available.

We also note that the intrabody screening step is readily scalable to the throughput of transient transfection as implemented by robotic liquid handling. This makes it ideal for interfacing with computational workflows such as Nb humanization and *de novo* design^34–36^ where many candidate Nb sequences are generated and must be screened intracellularly, as recently demonstrated using a split luciferase system.^37^ Moreover, this workflow should support development of intrabodies based on binding proteins other than Nbs, such as computationally-designed mini-binders^34^ and single-chain variable fragments (scFvs)^35^. Thus, our nanoBRET-based screening approach seems well suited to experimental validation of intracellular affinity reagents that are designed *in silico*, whether they are Nbs or other types of binding proteins.

## Methods

### Nb Library Biopanning

Three rounds of biopanning were conducted using a synthetic Nb phage-display library and established methodology as previously described^25^. Briefly, ITCH HECT domain (Sino Biologicals,11131-HNCE) was immobilized in a 96-well Immulon HBX microplate (ThermoFisher, 3855). For the first round of biopanning, wells were coated at 50μL well^-1^ with 0.06mg mL^-1^ ITCH in immobilization buffer (IB) comprised of 100mM NaHCO_3_ pH 8.3 and 150mM NaCl. For the subsequent rounds, wells were coated at 0.04mg mL^-1^ in IB overnight at 4°C. After plate coating, wells were then washed five times with 300μL of PBS-T (0.05% (v/v) Tween-20) and blocked for 2 hours with 300μL protein free blocking buffer (ThermoFisher, 37572). Post-blocking, wells were washed and 200μL of phage at 2.5×10^12^ CFU mL^-1^ was added and incubated at ambient conditions with shaking at 400rpm for 2 hours. Rounds two and three were performed similarly using 150μL of 6.67 x 10^10^ CFU mL^-1^ phage solution. After the addition of phage, the wells were washed multiple times with 300μL of PBS-T with increasing concentrations of Tween-20. In round 1, this varied from 0.1% to 0.5% (v/v) and by round 3 this was from 0.1% to 1% (v/v), generally in steps of 0.1% per wash. After washing, bound phage was cleaved in trypsin buffer (10mM Tris-HCL pH 7.4, 137mM NaCl, 1 mM CaCl_2_, 0.1 mg mL^-1^ trypsin). To deplete non-specific phage clones, the blocking buffer was varied in rounds two and three to 1% (w/v) BSA and 2% non-fat milk in PBST respectively. Outgrowth and purification of harvested phage round-to-round was done as described previously for this library.^25^

### Next Generation Sequencing (NGS) Analysis of Biopanning Outputs

Nb phagemids were purified from harvested phage as previously described^25^, and Nb sequences represented therein were identified and enumerated *via* NGS analysis as follows.

A previously described two-stage PCR approach^38^ was modified for use in generating an amplicon library for sequencing our Nb-encoding cassettes using Illumina instrumentation. The PCR primer sequences are listed in **Supplemental Table 1**.

In the first stage of PCR (PCR1) the forward (sense) primer (PCR1-Fwd) was used to incorporate the 3’ end of the Illumina universal library amplification forward primer (Illumina F seq) binding sequence, as well as a random nucleotide sequence of variable length [a universal molecular identifier (UMI) that also introduced stagger to support cluster recognition], at the 5’ end of the amplicon. The PCR1 reverse (antisense) primer (PCR1-Rev) was used to incorporate the 3’ end of the Illumina universal library amplification reverse primer (Ilumina R seq) binding sequence, as well as an index sequence of variable length (a barcode to enable multiplexing of amplicon libraries), at the 3’ end of the amplicon. Each PCR1 reaction (50 µL total) consisted of 200 ng of Nb phagemid DNA, 500 nM of PCR1-Fwd primers (eight variants mixed at equal molar ratio), 500 nM of PCR1-Rev primers (eight variants mixed at equal molar ratio), 0.5 units of Phusion High-Fidelity DNA Polymerase (New England Biolabs, M0530) with 1X buffer, 0.5 units of Deep Vent DNA Polymerase with 1X buffer (New England Biolabs, M0258), and 200 nM dNTPs (each mixed at equal molar ratio). Each Nb phagemid DNA sample was used to generate six identical PCR1 reactions. PCR1 cycling conditions were: 95`°` C for 1 min (1 cycle); 95`°` C for 15 sec, 65`°` C for 15 sec, 72`°` C for 20 sec (5 cycles); 72`°` C for 1 min (1 cycle). The six identical PCR1 reactions were then combined and their products purified using the NucleoSpin Gel and PCR Clean-Up kit (Macherey Nagel, 740609), eluting with 66 µL of Buffer NE to yield ∼60 µL of eluate.

In the second stage of PCR (PCR2) the forward (sense) (PCR2-Fwd) primer was used to incorporate the Illumina P5 adapter sequence at the 5’ end of the PCR1 products, and the reverse (antisense) (PCR2-Rev) primer was used to incorporate the Ilumina P7 adapter sequence at the 3’ end of the PCR1 products; addition of these adapters enables binding of the PCR2 products to the flow cell surface. Each PCR2 reaction (50 µL total) consisted of 10 µL of PCR1 product, 500 nM of PCR2-Fwd primer, 500 nM of PCR2-Rev primer, 0.5 units of Phusion High-Fidelity DNA Polymerase with 1X buffer, 0.5 units of Deep Vent DNA Polymerase with 1X buffer, and 200 nM dNTPs (each mixed at equal molar ratio). Each PCR1 product was used to generate six identical PCR2 reactions. PCR2 cycling conditions were: 95`°` C for 1 min (1 cycle); 95`°` C for 15 sec, 65`°` C for 15 sec, 72`°` C for 20 sec (15 cycles); 72`°` C for 20 sec (1 cycle). The six identical PCR2 reactions were then combined, purified using the QIAquick PCR Purification Kit (Qiagen, 28104), and size-selected using open-tank gel electrophoresis (1.2% agarose) or BluePippin (Sage Science) (1.5% agarose) followed by purification using the NucleoSpin Gel and PCR Clean-Up kit, eluting with 25 µL of water to yield ∼20 µL of eluate.

The concentration of each PCR2 product was measured using the Qubit dsDNA High Sensitivity Assay kit (Thermo Fisher Scientific, Q332851). Then the PCR2 products from the three biopanning round outputs (each set bearing a unique barcode) were combined at equal concentration to generate the final library, which was analyzed using 300 bp paired-end Illumina NGS (10-20M reads/PCR2 product).

Adapter sequences were trimmed using fastp^39^ and a custom script was used to collapse UMIs, identify the sequences encoding the three complementarity determining regions (CDRs) of each Nb represented in the NGS data, and assess the relative abundance of each unique Nb sequence after normalizing for coverage depth.

### Molecular Cloning

Cloning of mammalian expression constructs was done using Golden Gate parts and syntax derived for the mammalian toolkit (MTK).^31^ Enzymes were obtained from New England Biolabs, and cycling conditions were used according to the manufacturer’s instructions. An exception to the MTK syntax was made for BxB1 AttP and AttB containing plasmids due to an internal BsaI restriction site. *E. coli* expression vectors were constructed *via* HiFi assembly (New England Biolabs, E2621L) using chemically synthesized DNA fragments (Twist and IDT). A table summarizing the key plasmids used in this study, their encoded protein sequences, and details regarding their use is provided (**Supplemental Table 2**).

### Cell Culture and Cell Line Generation

HEK293T cells were grown in DMEM (ThermoFisher, 21063029) supplemented with 10% FBS (ThermoFisher, A4736301) and penicillin-streptomycin (ThermoFisher, 15070063). Cells were maintained in 5% CO_2_ at 37°C with regular passaging using 0.25% (w/v) Trypsin-EDTA. BxB1 landing pad cell lines containing an icasp9-based kill marker were generated using low multiplicity of infection lentivirus transduction, following a previously detailed protocol and design^40^. Single-cell clones of the transduced lines were then recovered *via* single-cell sorting using a FACSymphony S6 (BD Biosciences). Clones expressing high levels of the landing pad emirfp670 marker were qualitatively identified for further use. Nb knock-in cell lines were then generated using BxB1 recombination *via* co-transfection of a BxB1 recombinase expression plasmid and a donor plasmid. The transfection was conducted using Fugene 4k at a ratio of 60ng BxB1 plasmid and 930ng of donor diluted in serum-free media. Polyplexes were prepared with the addition of Fugene 4k to 4%(v/v) and incubation for 20 minutes. 100 μL of prepared polyplexes were added to 350,000 HEK293T cells in a 6-well plate. The donor plasmid contains a Tet-On3G doxycycline sensitive transcription factor (Takara) in frame with an upstream BxB1-AttB sequence. A second expression cassette in the same donor plasmid carries a dox-inducible promoter driving the Nb-mNG coding sequence with an internal ribosomal entry site (IRES) for polycistronic expression of a Zeocin (ThermoFisher, R25001) marker. Four days after transfection, negative selection was performed by the addition of AP1903 in DMSO to 10nM for 6 hours before a medium exchange into non-selective DMEM growth medium supplemented with 500ng/mL doxycycline-hyclate (Sigma-Aldrich, D9891). The following day, Zeocin was added to 300μg/mL and selection was kept through regular culture and expansion of the generated cell line until purity was established, about a week later.

### nanoBRET assay

HEK293T cells were seeded in optically-clear flat-bottom fibronectin-coated 384-well plates (Revvity, 6057600) in 50 μL of phenol-free DMEM growth medium at a density of 12,000 cells well^-1^. After overnight attachment, each well was transfected with 6 μL of polyplexes formed using DNA and Fugene 4K (Promega, E5911) following the manufacturer’s recommendations. Briefly, polyplexes were formed at a ratio of 1000 ng DNA per 200 μL of serum-free DMEM medium supplemented with Fugene 4K to 2% (v/v). This amounted to 30 ng of total plasmid per well in the 384-well plate. Cell seeding and transfection were carried out using a robotic liquid handler (Tecan Fluent). To fix the input mass of DNA for polyplex formation where the ratios of donor and acceptor plasmids were varied, a mass equivalent of pUC19 (NEB, N3041L) was substituted. For greater clarity, formulations of plasmids used for BRET transfection experiments in this study are given in **Supplemental Table 3**. BRET measurements were then carried out the next day with the per-well addition of 12 μL of Nano-Glo reagent (Promega, N2012) diluted 20-fold in complete medium without serum. The resulting luminescence was measured using a plate reader (Tecan Spark). A band filter of 415-485 nm was collected for donor luminescence and 505-650 nm was collected for acceptor luminescence. Where computed, the raw BRET and net BRET signals were calculated as previously described^41^ using an equation reproduced below, where is the measured luminescent flux at the given wavelength and C_f_ is a correction factor for spectral bleed between the acceptor and donor.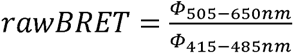;

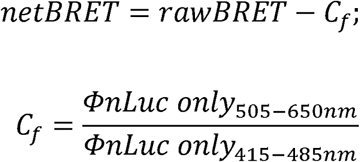

Acceptor (mNG) signal was measured using a Zeiss LSM900 confocal microscope equipped with an automated stage, using a 5x objective and 488 nm laser, and collecting 410nm-617nm. In optimization experiments we observed that transfecting more than 18 ng well^-1^ of mNG with non-interacting nLuc would result in unexpectedly high nanoBRET signal at or above 3 standard deviations above mean (**Supplemental Figure 9**), and so those data were not included for fitting nanoBRET titration data.

### Confocal Imaging

Live HEK293T cells were imaged in 10mM HEPES supplemented phenol-free complete DMEM using a Zeiss LSM900 with heated automated stage. mScarlet-I was excited at 561nm and emission collected at 536-700nm. mNeonGreen (mNG) was excited at 488nm and collected at 400-525nm. Hoechst 33342 (ThermoFisher, H3570), added to a final concentration of 1μg/mL, was imaged using 405nm excitation and 410-536nm emission.

### Recombinant Nb Production

Nbs were purified from *E. coli* using a C-terminal 6xHis tag and expressed using a pelB secretion signal driven, IPTG inducible periplasmic expression system following a previously described protocol.^42^ Briefly, BL21(DE3) cells bearing Nb expression constructs were grown in 3mL LB cultures at 37°C with shaking (300 rpm) for 16 h, then diluted 1:100 into terrific broth supplemented with 100μg/mL carbenicillin and further grown at 37°C. When the cultures reached an OD_600_ of 0.8-1.0 they were briefly chilled on ice to reach 25°C, IPTG was added to a final concentration of 1mM, and Nb expression was induced at 25°C with shaking (300 rpm) for 20 h. The following day, induced cultures were pelleted *via* centrifugation and weighed. For each gram of wet cell mass, 3mL of Tris-EDTA-sucrose (TES) buffer (200mM Tris-HCl pH 8.0, 0.5mM EDTA, 500mM sucrose) supplemented with protease inhibitor cocktail (ThermoFisher, A32963) were used to resuspend the cell pellet into a slurry. The slurry was incubated at 25°C for 45 min before addition of ice-cold water in a volume equal to the added TES buffer and further incubation on ice for 30 min with intermittent mixing by inversion. The resulting extract was then supplemented to 150mM NaCl, 2mM MgCl_2_, and 40mM imidazole before centrifugation at 20,000 xg for 15 min. The Nbs were then purified by recovery onto Ni-NTA (ThermoFisher, 88221) followed by elution using Tris-buffered saline (TBS) (20mM Tris-HCl pH 7.5, 150mM NaCl) supplemented with 500mM imidazole. The eluates were desalted into TBS *via* dialysis (ThermoFisher, 88400) overnight with two buffer exchanges.

### Biolayer Interferometry (BLI)

Binding of purified Nbs to their target was measured using an Octet biolayer interferometer (Sartorius) equipped with HIS1k biosensors (Sartorius, 18-5120). In general, 6xHis-tagged Nbs were loaded onto sensors at a concentration of 62.5nM when a concentration could be determined, and variable concentrations of ligand were used in the association step. All BLI measurements were conducted using a buffer comprised of 1x HBS+EP (Cytiva, BR100669) supplemented with 0.1% (w/v) bovine serum albumin and 0.6M sucrose to suppress non-specific binding.^43^

### In Vitro Enzymatic Assays

ITCH auto-ubiquitination was measured using endpoint TR-FRET (UBPbio, T4010) according to the manufacturer’s directions in a 384-well plate (Corning, 4514). Briefly, full-length ITCH was incubated with an E2 enzyme, E1 ligase, and a mixture of ubiquitin conjugated with either a terbium donor or fluorescein acceptor. The reaction was then charged with ATP and auto-ubiquitination of ITCH was measured by TR-FRET using a plate reader (Tecan Spark Cyto) with the reaction proceeding at 25`°`C. An excitation of 340nm/20 was used and the acceptor and donor filter settings were 490nm/20 and 520nm/20, respectively, with a time-delay and integration of 100μs and 400μs, respectively. TR-FRET signal was computed according to the manufacturer’s instructions and reported as values normalized to a control reaction. In TR-FRET inhibitor studies, Nbs were supplemented in the reaction at 200nM final concentration (2:1 Nb:ITCH molar excess), and clomipramine (MCE, HY-B0457) was supplemented to 250μM.

*In vitro* ITCH ubiquitination of LATS1 was studied in a enzyme reaction mix containing 0.6 μM ITCH (UBPbio, K1400), 1μM FLAG-LATS1 (Active Motif, 81209), 2uM UbE1 (UBPbio, E1100), 8uM UbE2D4 (UBPbio, C1700), 75uM HA-Ub (UBPbio, E1400) in 50mM Tris-HCl pH 8, 50mM NaCl, and 2.5mM MgCl_2_. Where specified, 1.8 μM Nb was supplemented, and ATP was added to 2mM final concentration before incubation at 37°C for 90 min. At the end of incubation, the reaction was quenched by heating to 98°C for 10 min in 1x Laemmli buffer supplemented with 5% (v/v) β-mercaptoethanol. The resulting *in vitro* reactions were then subjected to Western analysis following manufacturer’s instructions (BioRad) and probed using an AlexaFluor (AF) 647 conjugated anti-HA antibody (ThermoFisher, 26183A647) (1:1000 dilution) and an anti-FLAG-AF488 antibody (ThermoFisher, MA1-142-A488) (1:1000 dilution).

### Immunoassays and Western Analysis

For HA-ITCH immunoprecipitation (IP) experiments, HEK293T cells were seeded in a 12-well plate at a density of 50,000 cells per well and allowed to attach overnight. They were then transfected with a Fugene 4K polyplex mixture of 100ng HA-ITCH plasmid, 200ng of myc-Ub plasmid, and 50ng of either Nb66-mNG, α-Histone-mNG, or pUC19, with the remaining balance made with pUC19 to a total mass of 1000ng plasmid per well. At 20 h post-transfection the proteasome inhibitor MG132 (Sigma-Aldrich, M7449) was added to 5μM and the cells incubated at 37°C for 5 h. The cells were then lysed in an IP buffer containing 10 mM Tris-HCl pH 7.5, 150 mM NaCl, 0.5 mM EDTA, 0.5 %(v/v) NP-40 supplemented with protease inhibitor cocktail (ThermoFisher, 78440). Lysates were incubated with anti-HA magnetic beads (Sigma-Aldrich, SAE1097) at 25°C for 1 h and elution was performed by heating at 98°C for 5 min in 2x Laemmli buffer. The eluate was then aspirated from the beads and β-mercaptoethanol was added to 5% (v/v) before heating again at 98°C for 10 min.

For mNG IP experiments, HEK293T cells were seeded at 500,000 cells well^-1^ in a 6-well plate before transfection the following morning using Fugene 4k polyplexes containing 1800ng of HA-ITCH plasmid and 200ng of either mNG, Nb66-mNG, or α-Histone-mNG plasmid. At 20 h post-transfection the cells were harvested in lysis buffer and incubated with anti-mNeonGreen magnetic agarose (Proteintech, ntmak-20) at 25°C for 1 h. Elution was conducted by heating the beads in 2x Laemmli buffer at 98°C for 5 min. The eluate was then spiked with 5% (v/v) β-mercaptoethanol and incubated at 98°C for an additional 10 min.

Lysates were size-fractionated *via* SDS-PAGE using a 4-15% gradient gel (Bio-Rad, 4561086) with a fluorescent molecular weight marker (Bio-Rad, 1610376), and proteins were transferred to a low-fluorescence PVDF membrane (Bio-Rad, 1620260). The membrane was incubated in a commercial blocking buffer (Bio-Rad, 12010020). Primary antibody incubations were done overnight at 4°C and secondary antibody incubations were done for 2 h at 4°C.

For intracellular ITCH auto-ubiquitination studies, a primary antibody cocktail of anti-HA-HRP (CST, 2999S) (1:1000), anti-GAPDH-AF555 (ThermoFisher, MA5 15738 A55) (1:2000), and anti-mycTag (Sigma-Aldrich, pla0001) (1:1000) in blocking buffer was used. The secondary antibody cocktail contained anti-rabbit AF800 (ThermoFisher,A32735) (1:1000) in blocking buffer supplemented with 0.05%(w/v) sodium dodecyl sulfate (SDS).

For mNG Western blots, the primary antibody cocktail contained anti-HA-AF647 (CST, 3444) (1:1000), anti-GAPDH (CST, 2118) (1:2000), and anti-mNG (Proteintech, 32F6) (1:1000). The secondary antibody cocktail contained anti-rabbit IgG-AF800 (ThermoFisher, A32735) (1:1000) and anti-mouse IgG-HRP (CST, 7076) (1:10,000) in blocking buffer supplemented with 0.05%(w/v) SDS. HRP was detected using ECL chemiluminescent substrate (ThermoFisher, 34580).

### Data Analysis and Statistics

Plate-reader data were aggregated using custom scripts written in Python 3.11, and visualized in GraphPad Prism 11. Where performed, statistical analysis was done using GraphPad Prism 11 built-in functions. Qualitative image data were acquired using Zeiss Zen Blue Ver3.8, and quantitative data were acquired using CellProfiler 3.8 prior to import into GraphPad Prism 11. Equilibrium model fitting was done using a single site equilibrium model:

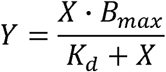

The average fit parameters and their 95% confidence intervals were found through bootstrapping with 500 iterations.

## Acknowledgements

We thank Catherine M. Mageeney (SNL) for critical review of the manuscript. Sandia National Laboratories is a multimission laboratory managed and operated by National Technology & Engineering Solutions of Sandia, LLC, a wholly owned subsidiary of Honeywell International Inc., for the U.S. Department of Energy’s National Nuclear Security Administration under contract DE-NA0003525. This paper describes objective technical results and analysis. Any subjective views or opinions that might be expressed in the paper do not necessarily represent the views of the U.S. Department of Energy or the United States Government.

