## supplemental figures for "Intracellular screening of nanobodies reveals an intrabody that inhibits ITCH E3 ubiquitin ligase"

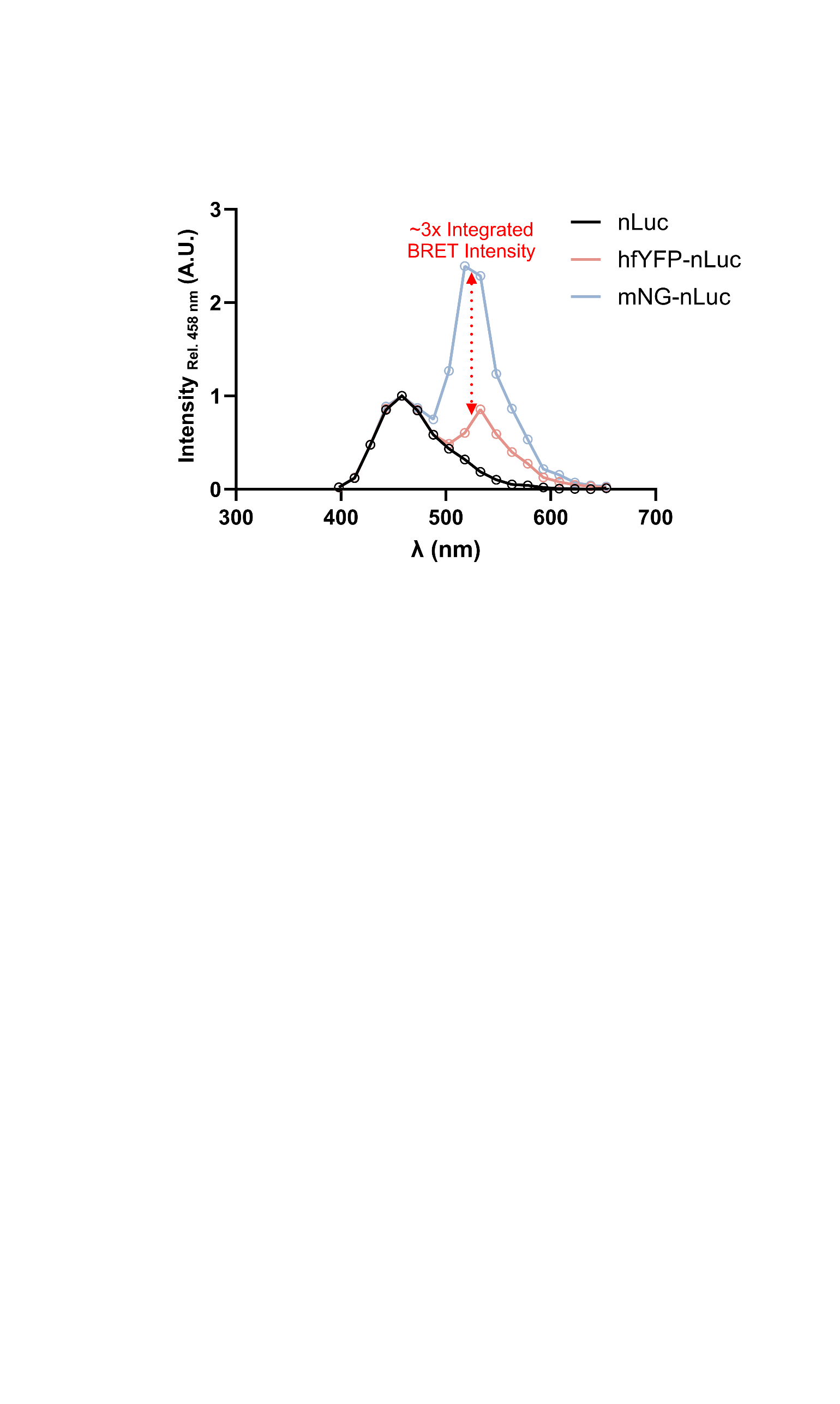

**Supplemental Figure 1.** **Higher BRET efficiency when using mNG, rather than hfYFP, as the acceptor.** HEK293T cells were transfected with either hfYFP-nLuc or mNG-nLuc, and luminescence spectra were acquired. A simple Riemann sum between 503-653 nm was computed to determine the integrated BRET intensity.

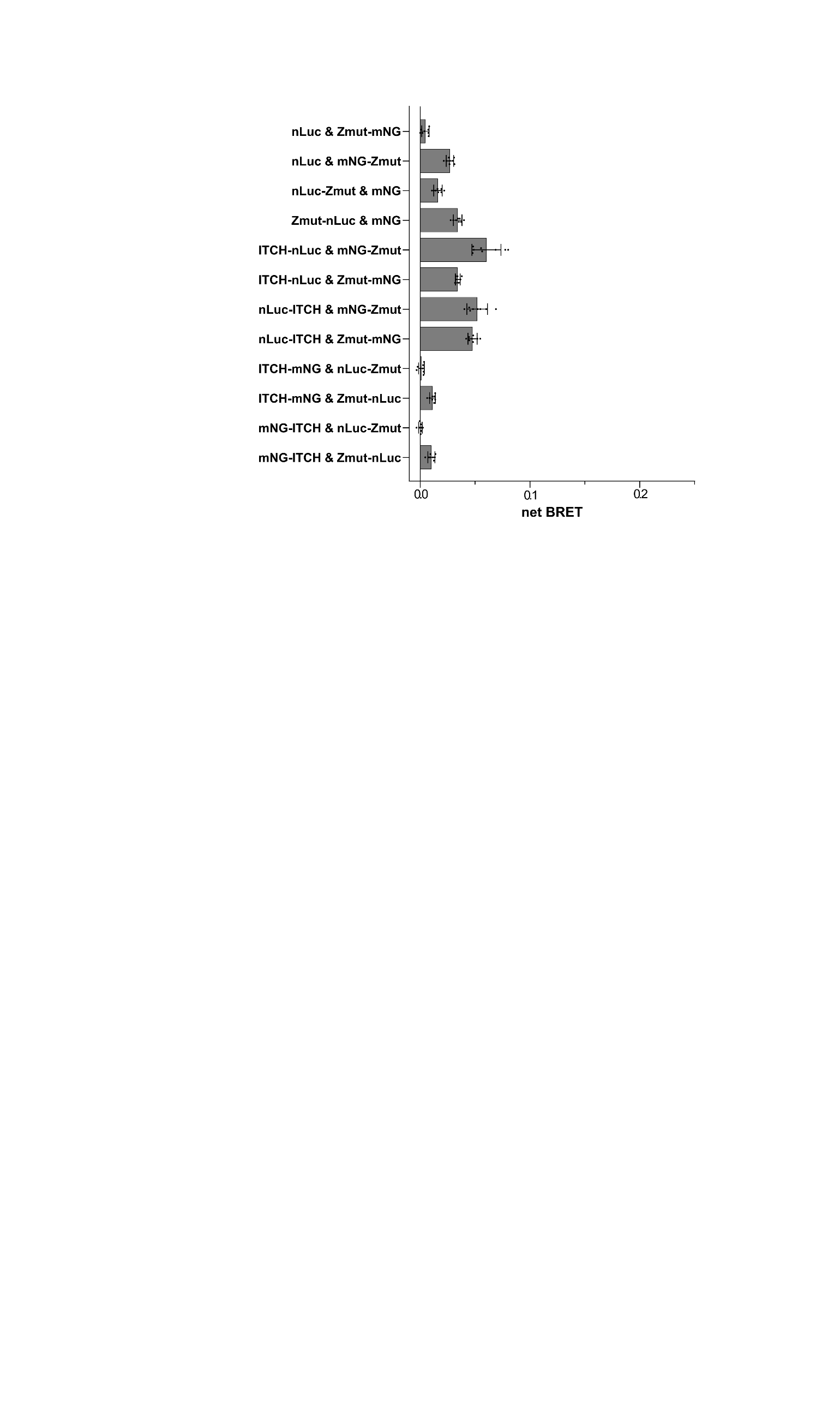

**Supplemental Figure 2.** **Combinatorial N- and C- terminus tagging optimization for an ITCH + Z_mut_ PPI nanoBRET assay.** Luminescence was measured at 24 h post-transfection. Net BRET signal was computed for each condition (n=8, replicate wells).

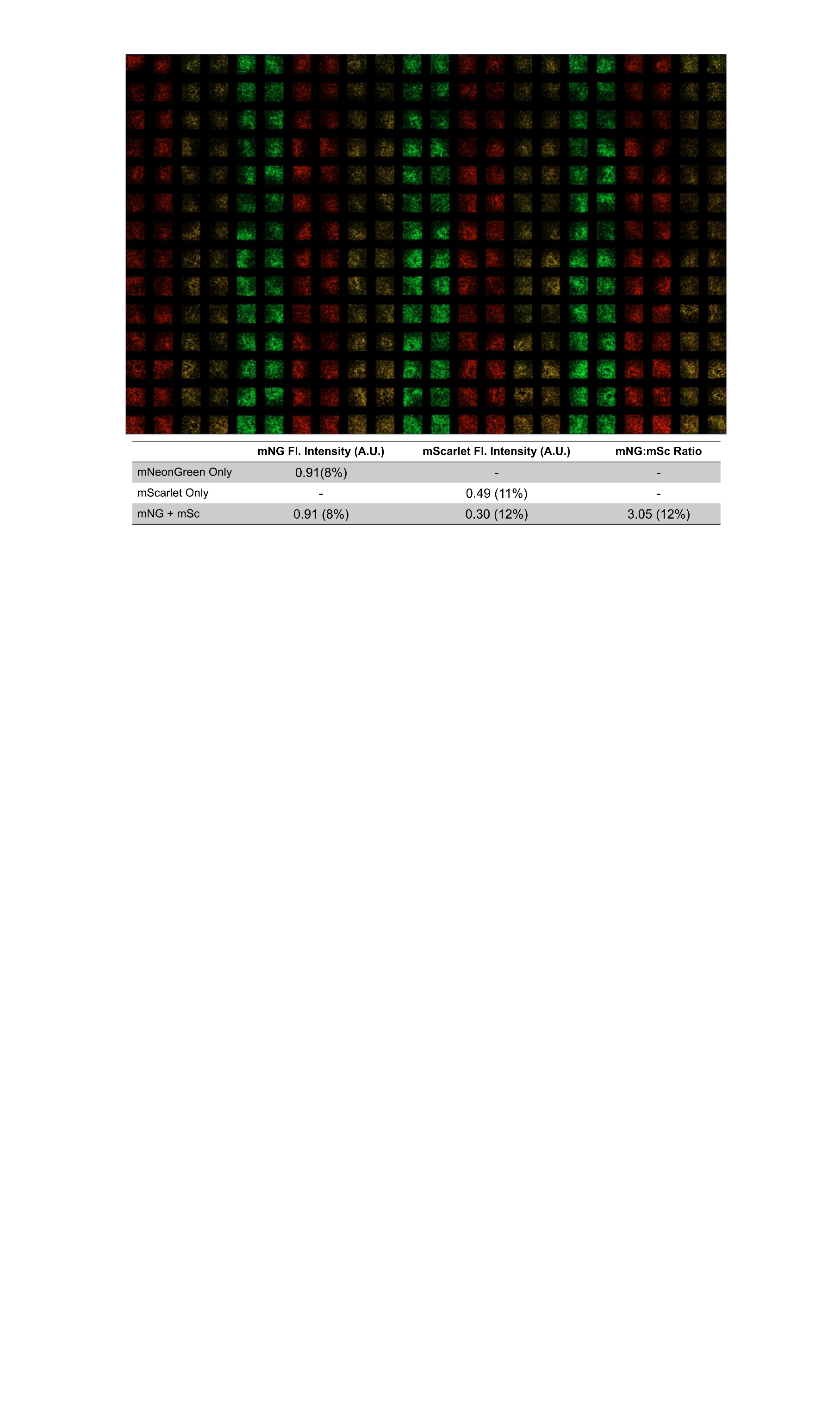

**Supplemental Figure 3. Representative robotic liquid handler transfection benchmark.** A 384-well plate was seeded with HEK293T cells, and the cells were transfected in quadruplicate sets using either an mNG (green pseudocolor) plasmid, mScarlet (red pseudocolor) plasmid, or both. For these experiments the cells in the outer rim of the plate were not transfected. Fluorescence intensity was quantified and normalized to the number of cells *per* well as estimated using a nuclear stain. The table summarizes the mean values across each condition, with the coefficient of variation for that condition and measurement in parentheses.

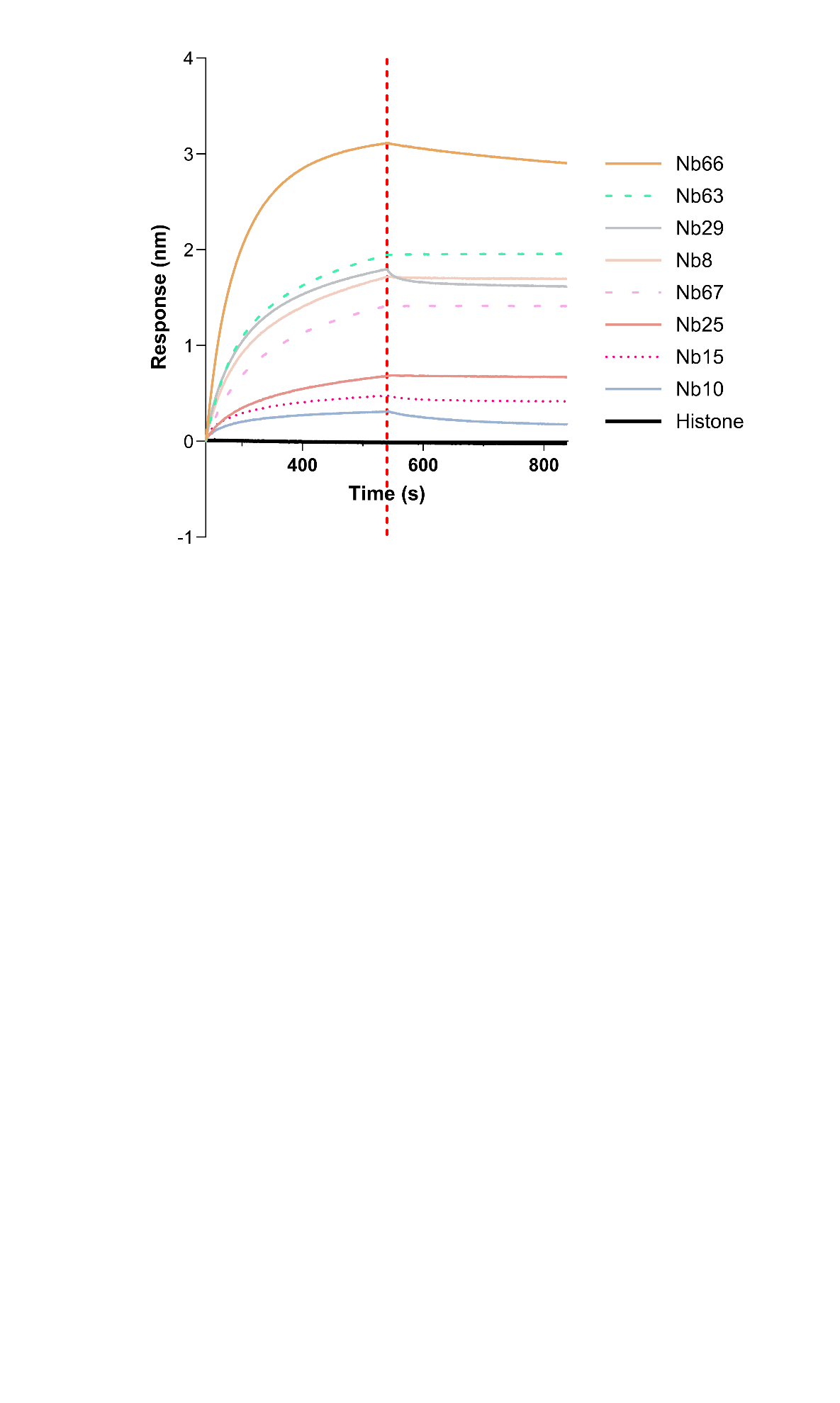

**Supplemental Figure 4. α-ITCH intrabodies bind to the ITCH HECT domain *in vitro*.** The BLI sensor was loaded with Nb and then pulsed with ITCH HECT domain (250 nM) for 540 seconds. The dashed red line indicates the transition from Nb-ITCH association to dissociation.

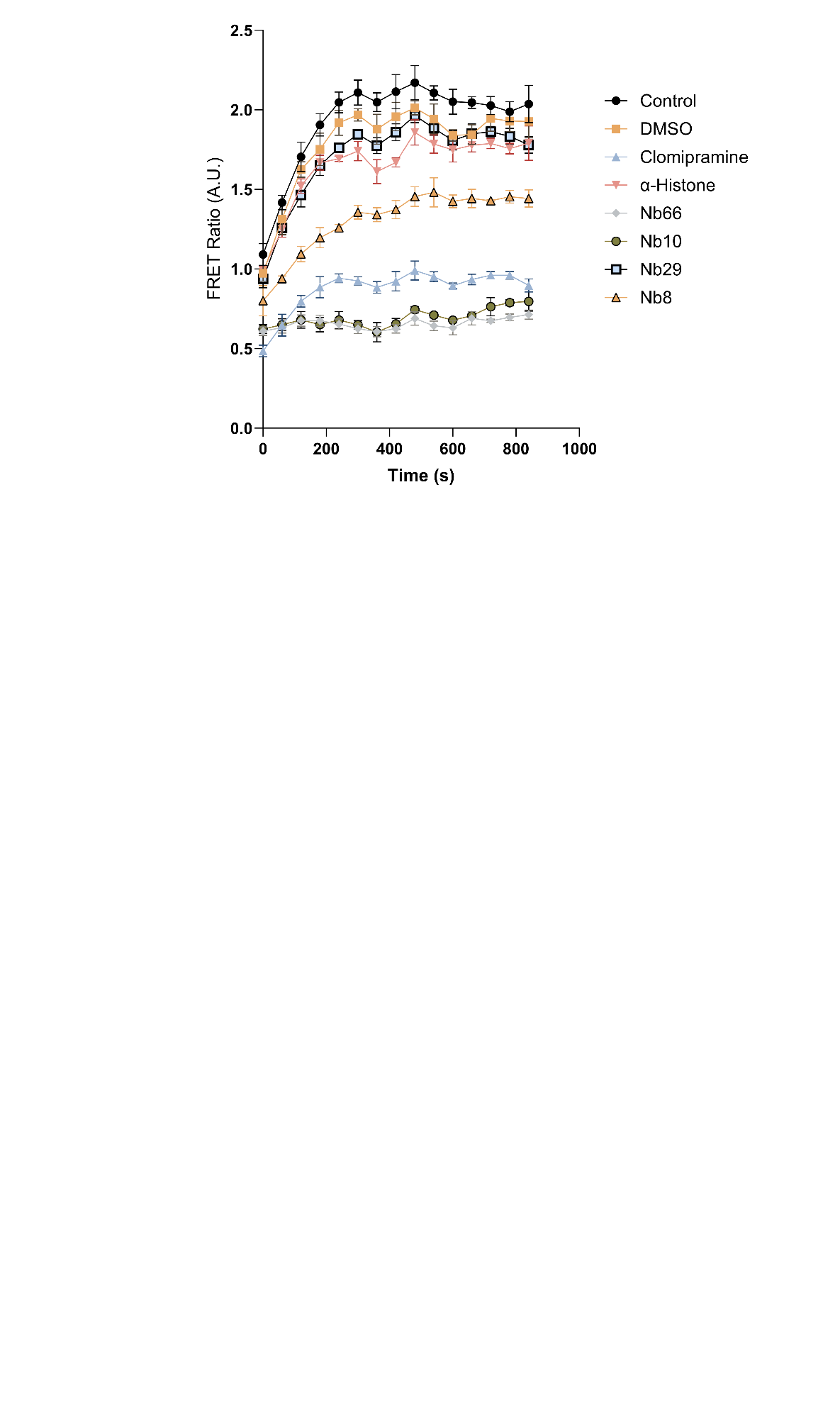

**Supplemental Figure 5.** **α-ITCH intrabodies inhibit ITCH auto-ubiquitination activity *in vitro*.** Each Nb (200 nM) was added to the ITCH HECT domain (100 nm) to achieve two-fold molar excess, and ITCH auto-ubiquitination was measured using an end-point TR-FRET assay (n=4 replicate wells). In positive control reactions, clomipramine (a small-molecule inhibitor of ITCH) was added (250 μM) instead of a Nb; and in negative control reactions only buffered saline was added. TR-FRET signal from each reaction was normalized to a photobleaching control and then to the average of the positive control reactions.

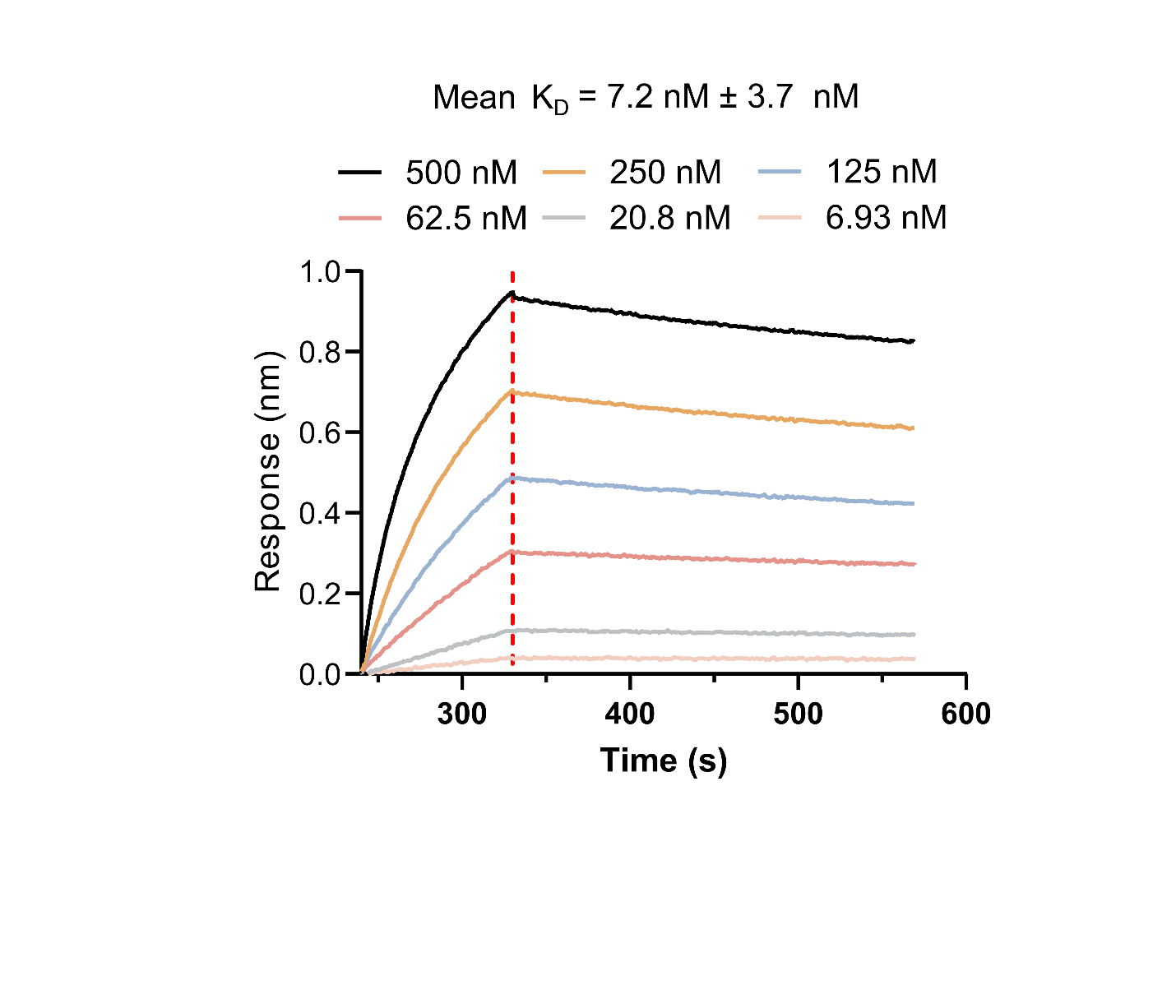

**Supplemental Figure 6. Measurement of Nb66 affinity for the ITCH HECT domain *in vitro*.** The BLI sensor was loaded with α-ITCH intrabody Nb66 and then pulsed with the ITCH HECT domain at the six specified concentrations (range: 6.93-500 nM) for 330 seconds. The red dashed line indicates the transition from Nb-ITCH association to dissociation. The mean and standard deviation of the six fitted K_d_s are provided.

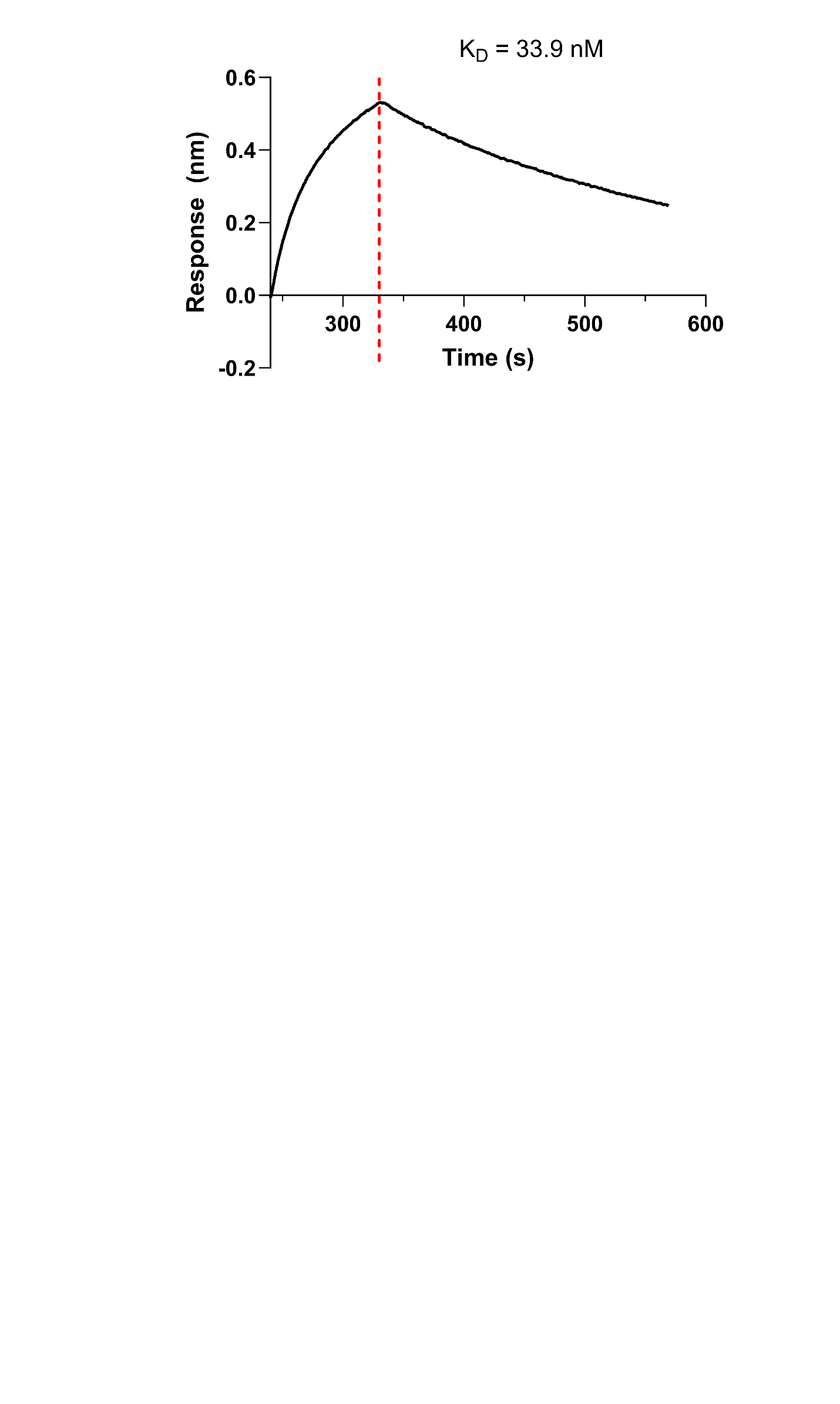

**Supplemental Figure 7. Nb66 binds to full-length ITCH *in vitro*.** The BLI sensor was loaded with 62.5 nM α-ITCH intrabody Nb66 and then pulsed with full-length ITCH (250 nM) for 330 seconds. The dashed red line indicates the transition from Nb-ITCH association to dissociation. The calculated K_d_ value is indicated.

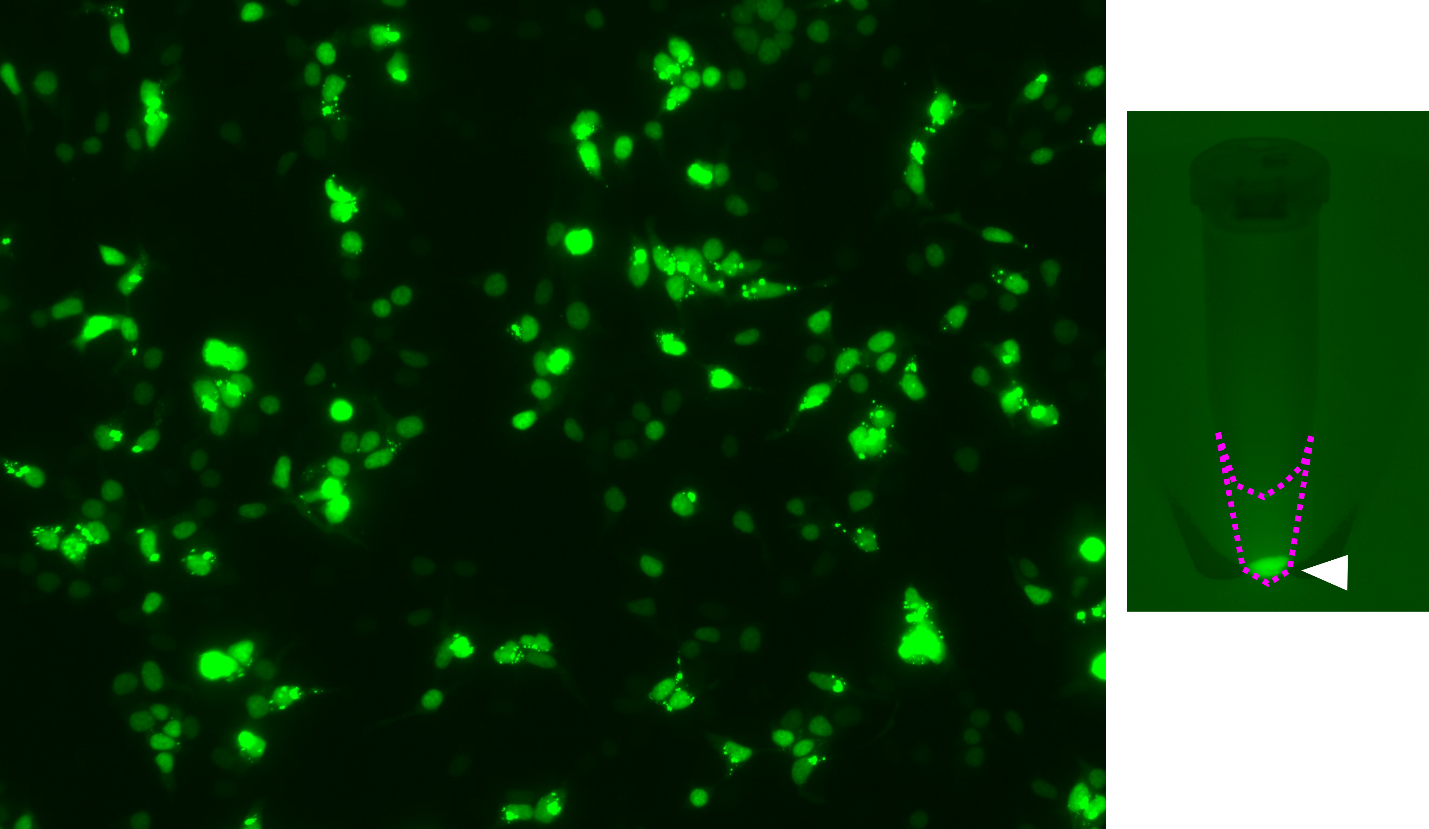

**Supplemental Figure 8. α-Histone-mNG intrabody in cells and cell lysates.** HEK293T cells were transfected with an expression construct for production of α-Histone-mNG. At 24 h post-transfection the cells were imaged using widefield microscopy with a GFP fluorescence excitation/emission filter set (left). Nuclear localization of α-Histone-mNG indicates proper function of this histone-targeted intrabody. Lysates from the transfected cells were fractionated *via* centrifugation (18,000 *x g* for 10 min), and α-Histone-mNG was visualized using blue illumination in combination with a GFP emission filter (right). The dotted magenta line demarcates the aqueous phase of the lysate; the white arrowhead points to the insoluble fraction of the lysate, which apparently contains most of the α-Histone-mNG as indicated by its fluorescence intensity.

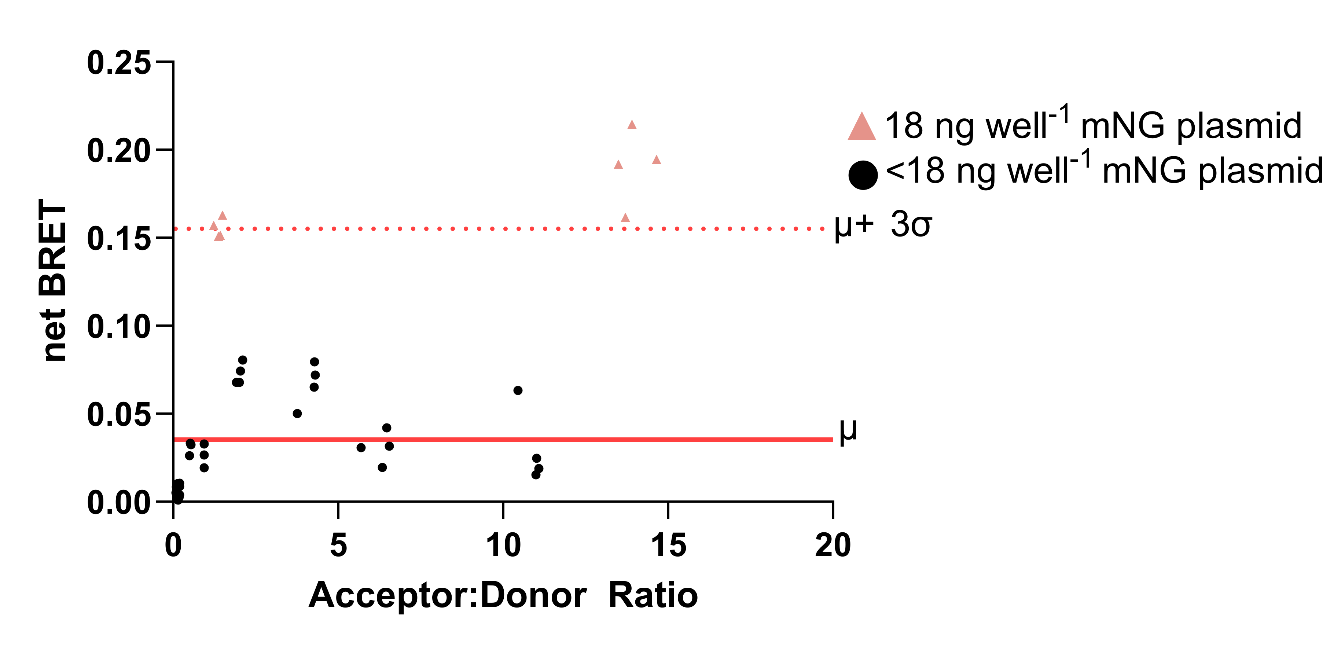

**Supplemental Figure 9.** nanoBRET titration assay performed in HEK293T cells transfected with varying concentrations of nLuc and mNG. The net BRET of each well (y-axis) is plotted against the measured Acceptor:Donor Ratio (x-axis). The mean, μ, of the net BRET is given by the solid red line and the mean plus three standard deviations, 3σ, by a dotted red line.
